# Monitoring Forest Foliar Moisture Using Sentinel-2 Reflectance and Radiative Transfer Model Inversion

**DOI:** 10.1101/2025.06.25.661444

**Authors:** Ivan Kotzur, Matthias Boer, Marta Yebra, Ben D Moore

**Affiliations:** This project has been assisted by the New South Wales Government Department of Climate Change, Energy, The Environment and Water. This work was also assisted by the National Computation Infrastructure through the Adapter Allocation Scheme

## Abstract

Foliar moisture content (FMC) of forests canopies is important for understanding biodiversity, including animal habitat quality for forest-dependent fauna and forest fire risk. In this study, we modelled spatiotemporal variation in FMC using optical remote sensing. We inverted the PROSPECT and GEOSAIL Radiative Transfer Models (RTM) on Sentinel-2 satellite reflectance data of 20 m ground resolution to retrieve FMC, and then predicted FMC at sub-continental scales using a random forest (RF) regression emulation of the RTM model. An RTM look-up-table, from earlier work, was filtered by ecological criteria from 24 sites sampled along precipitation and canopy cover gradients and was inverted by using the spectral angle, as a merit function. The RF emulator allowed efficient computation and presentation of an extensive and fine-scale forest FMC data cube of interest in animal, fire and plant ecology. Our RTM-based, predictions of forest and woodland FMC had a root mean square error (RMSE) of 19.9% of dry matter content and explained more than 60% of variance (r^2^ = 0.62). This represents an improvement over previous models using reflectance data of coarser spatial resolution, particularly in terms of explained variance (r^2^=0.17 and RMSE=32%). The emulator model achieved similar performance to the RTM with slightly larger error (RMSE = 21.77%) and smaller explained variance (r^2^ = 0.54). Overall, both models performed best in forests, and woodlands of moderate to high canopy density (> ∼0.75 LAI). This study demonstrates that FMC can be monitored at spatial resolutions that allow intra- or inter-landscape patterns to be resolved surpassing previous capabilities.

**Index Terms:** Foliar moisture content, FMC, remote sensing, radiative-transfer, forest, inversion, Sentinel-2, emulator, koala

## I. Introduction

CLIMATE and weather are important drivers of the water status of forested ecosystems, with great consequences for ecosystem structure and function. These include effects on plant species composition and animal habitat quality, to the regulation of water distribution and storage, and susceptibility to disturbances like windthrow, die-back and fire. In many biomes, periods of below-average rainfall are becoming more extreme and more frequent [1]. Coinciding with these droughts are increases in average air temperatures, and frequency and magnitude of positive temperature extremes [2]. Droughts and heatwaves can reduce plant productivity, alter their water use, and increase hydraulic failure and mortality [3], [4]. Animals may be impacted by such variations through changes to habitat quality including food quality and quantity, microclimate and water supply [5], [6]. Climate extremes also contribute to increased potential for large, high-intensity forest fires, which have become common in the 21^st^ century in Australia and other continents, due to forest fuels drying out, more often, over larger fractions of landscape [7], [8], [9].

Drying of vegetation reduces the water supply to many animals and limits their capacity to thermoregulate. Animals that rely on dietary pre-formed water, particularly mammalian arboreal herbivores, can usually meet their thermoregulatory water demands from the consumption of foliage alone [10], [11]. However, these demands increase along with exposure to high temperatures [12]. Furthermore, thermal stress for animals commonly coincides with drought stress on plants particularly where dry seasons are also warm seasons [13], meaning water supply declines as demand increases [14]. This challenge is encountered by all arboreal folivores, including primates, and koalas and possums in Australia [15], [16]. Thermo-physiological models of koala distribution recognise that foliar water content can constrain the species’ tolerance of heat stress, and that leaf moisture content varies at a fine scale over space and time [17]. Such models of koala distribution or similar species could be greatly improved if they were to be based on accurate predictions of spatiotemporal variation in foliage moisture at fine spatial and temporal resolution. This would enable the identification of refugial habitat under projected future climates [17]. Other animals are impacted by declines in vegetation moisture including many insects that need to avoid desiccation during periods of high temperatures by consuming water, or which rely on turgor pressure to facilitate feeding on phloem or xylem, such as aphids [18], [19]. These impacts are propagated throughout canopy food webs, as arthropods support many higher trophic levels, including other invertebrates and vertebrates [20].

The main drivers of variation in foliar moisture content (FMC) at the regional scales include, positively, rainfall and soil moisture, and negatively, atmospheric moisture demand [21]. Important drivers of variation in FMC at finer scales include soil moisture, topographical position, aspect, wind exposure, insolation and the drought tolerance of tree species [22]. Variation in prevailing leaf moisture content of the eucalypt forests and woodlands which are common in Australia, appears to be large. For instance, in the tropical north-east of Queensland, forests of *Eucalyptus cambageana* can have leaf water contents of up to 223% of dry weight, while it can be as low as 75% in *Eucalyptus brownii* further west on the central, semi-arid plains [23], [24]. In a similar, nearby ecosystem, the seasonal FMC range within species maybe as much as 50% of the leaf dry weight [25].

Mapping spatiotemporal variation in FMC is also of great importance in the context of fire ecology and fire management [26], which has driven most of the research into FMC. Declines in the moisture content of foliage and small-diameter twigs (i.e. live fuel) increases the availability of fuel for combustion [27]. This decline in live fuel moisture content is correlated with an increase in the flammability of vegetation and in the cumulative area of landscapes burnt by fire [8], [28], [29]. The dryness of live fuels is particularly important in forest and woodland environments which are inherently fire prone due to the abundance of plant biomass that becomes available for combustion when it dries out to ignitable levels [27]. As live fuel moisture drops below thresholds, the likelihood of fire ignition, as well as high fire severity and rates of spread increase [8], [9], [29], [30]. The largest forest fires tend to occur following ecological droughts when the ability of trees to maintain hydraulic function has been compromised and the foliage moisture content is low, as seen in widespread instances of canopy browning or dieback [3], [7], [31]. In many forest regions in Australia and other continents droughts are projected to increase in frequency and duration with climate change, resulting in forest fuels being highly flammable more often and for longer periods [32], [33]. Future fire management will require predictions of foliar moisture content at spatiotemporal scales relevant to plant ecophysiology and operational fire management (e.g. within tree communities and amongst stands, respectively).

FMC is the ratio of water to dry matter of foliage in live vegetation:

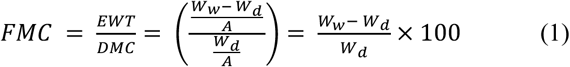

where *EWT* is equivalent water thickness or water mass per unit leaf area, *DMC* is dry matter content or the dry weight per unit leaf area, *W_w_* is fresh or wet weight, *W_d_* is dry weight and *A* is leaf area. Therefore, DMC and *W_d_* are equivalent, however we use them both in denoting units of FMC to indicate if whether the variable is predicted (DMC) or observed (*W_d_*, i.e. dry weight).

There are several approaches to FMC prediction using optical remote sensing. Two common approaches are empirical models, such as those based only on spectral vegetation indices [34], and radiative-transfer models (RTM). RTMs have been used to predict live fuel moisture content for fire behaviour modelling at scales from landscape to continental [28], [35]. This has been aided by the development of global databases of field measured fuel moisture content [36]. In RTM methodologies, leaf and canopy spectra may be inverted on remotely-sensed spectral data from vegetation to retrieve biochemical or biophysical variables coupled with the initially modelled spectra [37], [38]. Compared to empirically-derived remote sensing models (i.e. correlative models), this approach increases independence of the results from effects of vegetation type or condition and satellite sensor [39]. RTMs are physically-based models, based on leaf and canopy biophysical and biochemical parameters, that approximate the reflectance spectra of leaves or canopies. This approach allows for similar spectra to be produced by different sets of input parameters during inversion of the RTM (i.e. retrieval of a parameter from the spectra); this is referred to as the “ill-posed” problem [38]. The magnitude of this problem of indetermination can be reduced by eliminating most unrealistic spectral-parameter combinations by constraining the parameters to an observed, eco-physiological space. Jurdao et al. [40] used an RTM to predict FMC of woodlands in two Mediterranean ecosystems which included eucalypts, producing a look-up-table (LUT) to solve the inversion, and this has been applied successfully to Australian ecosystems [35]. However, the spatial resolution of the reflectance data used previously, from the Moderate-resolution Imaging Spectroradiometer (MODIS), is too coarse (MCD43A4, 500m) to resolve individual tree canopies, although significant variation in FMC occurs at that scale [33]. Valuable insights into water availability for fauna and refugia from drought, as well as drought impacts on vegetation and forest fuel dynamics, may be gained by producing finer-resolution FMC predictions at tree canopy scales.

Solar irradiation interacts most strongly with leaves in upper canopy layers. Interactions include the absorption of light by water in leaf mesophyll cells, particularly in the short-wave infrared wavelengths (1.3-2.6 mm, SWIR, [41], [42]. Across these wavelengths, dehydration results in a general increase in reflection due to reduced absorption and increased scattering and reflection from the enlarged interfaces between air and cells [43]. Two leaf water absorption features also occur in the near-infrared region of the electromagnetic spectrum (0.7-1.3 mm, NIR). Secondary effects of reduced leaf water content include browning at very low moisture levels, apparent in the visible spectrum (0.4-0.7 mm), due to relative changes in absorption by pigments [44]. The wavelengths at which reduced water content directly alter spectra are more universal among plant species than for pigments. The spectral changes brought about by the internal structure of leaves does not occur at discrete wavelengths but produces a broad spectral response, which is more subtle than that caused by water absorption features. In well-hydrated leaves, internal structure is masked by water absorption in mesophyll cells. These physical relations of light and moisture provide a pathway to quantify foliar water content [45]. FMC can be expressed as a water thickness measure or on a mass basis, the former being detectable in spectra, as relatively more light is attenuated in thicker columns of water [46]. However, this expression relies also on measurement of the leaf dry matter content which may also vary seasonally, and in response to drying of the foliage [47]. The conversion of FMC on a dry weight basis to water content (WC) on a fresh weight basis (i.e. total weight) produces a metric that is more intuitive.

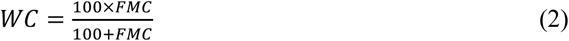

Where WC is water content of leaves or canopy on fresh weight basis, and FMC is that on a dry weight basis (Equation 1). Water content has been identified as an influential global leaf trait, particularly in predicting photosynthesis and leaf area [48].

The aim of this study was to accurately predict FMC at moderately fine spatial resolution (∼20 m) in forests and woodlands of south-east Australia. To this end, we inverted an RTM to retrieve FMC by first adapting the LUT of the existing RTM to Sentinel-2 specifications and applying specific ecological criteria of the study system to the LUT. This RTM LUT had previously been inverted only on MODIS reflectance data which has moderate spatial resolution.

## II. Methodology

We adapted existing RTM outputs by convolving the spectra to Sentinel-2 bands and filtering these RTM outputs based on our field observations of tree FMC (Fig. 1). This was followed by inversion of the RTM outputs on Sentinel-2 reflectance at our field observations (section B. Field Data), including inversion optimisation, and validation of these FMC predictions. We then built a dataset of FMC predictions with corresponding Sentinel-2 reflectance for random, forest pixels within the study area, to train and optimise a random forest (RF) model that efficiently emulated the initial RTM inversion for predicting FMC. We then used the RF model to predict FMC for the same field sites as earlier and validated this emulation model. Following this step, we applied the trained, emulator model to generate time series’ of spatially-explicit FMC predictions for all available Sentinel-2 reflectance data (2015 to mid-2024) in forests and woodlands across the study area as defined by a landcover mask that included pixels of native tree canopies.

**Fig. 1.**
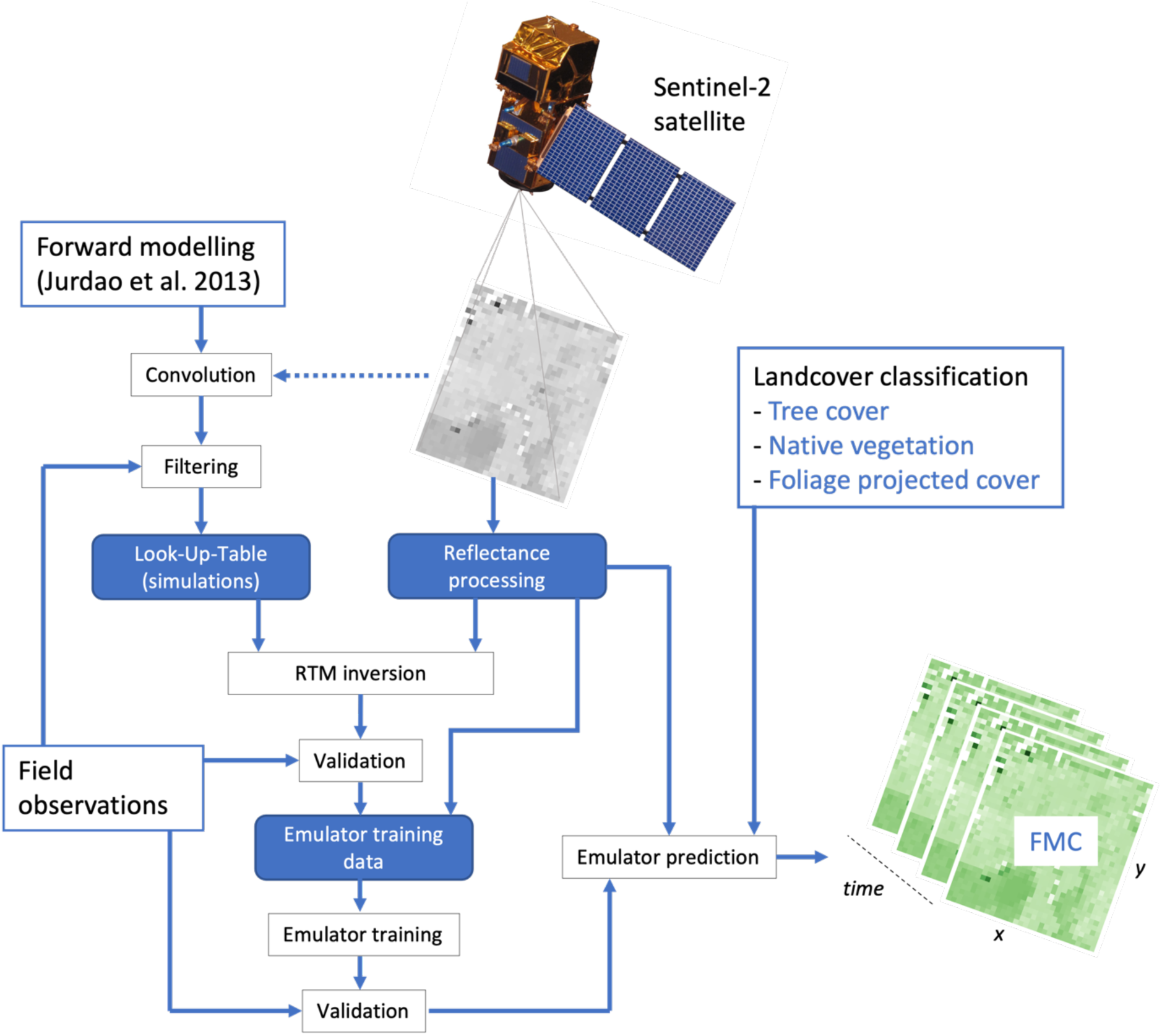
Methodology of RTM inversion and random forest emulator-based prediction of FMC in forest and woodlands.

### A. Study Area

The study area comprised forests and woodlands of continental New South Wales, a state in south-eastern Australia (Fig. 2). The area includes temperate, sub-tropical and alpine climates with cool winters and hot, dry summers in the centre and west, mild summers in the eastern ranges and hot humid summers on the north-eastern coastal fringe [49]. Treed ecosystems in the area mainly comprise dry-sclerophyll eucalypt forests and woodlands (the eucalypts include three genera, *Eucalyptus*, *Corymbia* and *Angophora*). The coastal fringe and mountains include wet sclerophyll forests and rainforest, and in the centre and west are eucalypt woodlands [50].

**Fig. 2.**
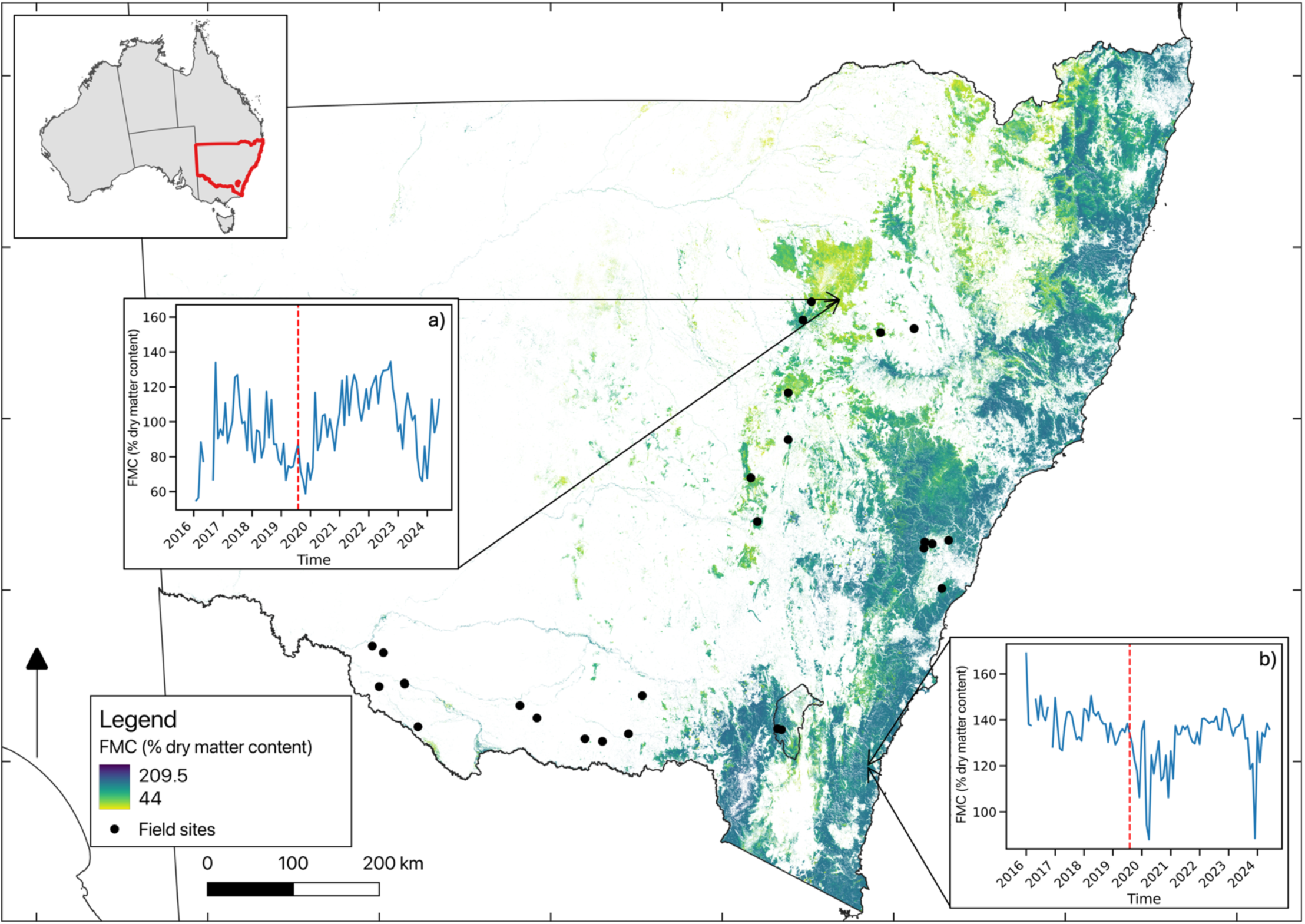
Foliar moisture content (FMC, % dry matter content) of forests and woodlands in NSW, Australia for July 2019 (monthly mean, calculated from the FMC data cube presented in this study). The inset timeseries show monthly means across 64 km^2^ areas (i.e. 8 km x 8 km) of the entire FMC data cube (2015-2024) in a) Pilliga Forest and b) Deua National Park. These two areas are ecologically significant - Pilliga supports koala populations vulnerable to heat and drought stress, while Deua National Park experienced severe bushfires during the summer of 2019/20. The red dashed line indicates the month displayed in the main map (i.e. July 2019).

### B. Field Data

Field observations of FMC were collected at 24 sites in different eucalypt-dominated forests and woodlands between 7/1/2022 and 21/03/2022, which was a relatively wet season in much of the study area (Fig. 2, Table S2, Table S3, Table S4). Field data collection was focused on environments with moderate to low moisture availability at the time, to capture a wide range of FMC values for the validation. For each sample, 20 g of leaves were cut from a small, sunlit branch, approximately 9 m off the ground, using a pole pruner, and sealed in an airtight, chilled, dark container. These fresh samples were weighed with a precision balance (±0.01 g) at the end of each day’s sampling, taken to the laboratory still refrigerated and then oven-dried. After 24 h drying at 105 °C samples were reweighed [51], and FMC was then calculated, according to Equation 1.

At each site, three samples were collected from the dominant tree canopies within the extent of a 20 x 20 m plot (i.e. one pixel). These plots typically included one or more trees, depending on the structure of the vegetation. We located the centre of the plot using a handheld GPS (model: Garmin ‘inReach Explorer+’, 3m-5m accuracy). We sampled each site at least twice, timing collections to occur on days without rainfall, within two days of a satellite overpass, to minimise temporal discrepancies between FMC and reflectance data. The field measurements of FMC were supplemented by previously published FMC observations from Namadgi National Park, ACT [36] (Fig. 2).

We undertook canopy cover photography at sites to measure leaf area index (LAI). Vertical, upward facing photographs were taken of the overstorey, five per site, using a digital single lens reflex camera with 50mm (35mm equivalent) lens and monopod, following [52], [53]. These images were later used to estimate LAI based on gap-fraction classification, using the software Digital Cover Photography Toolbox (version 3.13, [53]). The LAI results from the five images were averaged to produce a plot level LAI value.

### C. Radiative Transfer Modelling

The reflectance data used to invert the RTM in this study were sourced from Geoscience Australia’s Sentinel-2 Multispectral Instrument (MSI), Digital Earth Australia (DEA) Surface Reflectance datasets [54], [55]. The ground resolution of the reflectance data was 20 m. An existing LUT of simulated spectra of forest and woodland canopies [40] was compared to the observed reflectance data and the median FMC of a set of the most similar simulations was taken as the final prediction. During inversion, we performed an optimisation based on RMSE and r^2^ [35] of the set size (*n*=1 to *n*=750, i.e. model version M1), and of the set of size *n*=40 as per [40] (i.e. model version M2).

To account for errors in georeferenced satellite data and GPS data from field observations, we applied the RTM inversion to a window of 3 x 3 pixels around the central coordinate of field observations and took the mean of these nine FMC estimates as the site-level result. To make use of all possible field observations, while also limiting temporal differences in FMC and reflectance, we used satellite data if it had been collected within 10 days before or after each field sampling date and was flagged as good quality by the data custodians (i.e. not cloudy). We applied the inversion, and subsequent emulator model, to ‘tree cover’ pixels (i.e. forest and woodlands) from the NSW Native Vegetation Extent (v1p4) product, which is based on Satellite pour l’Observation de la Terre 5 (SPOT5) [56], upscaled using mode resampling [57].

We convolved the reflectance spectra of the existing LUT from continuous spectra spanning visible to SWIR wavelengths into the bands of the Sentinel-2 sensor (Table S1), following equation 5:

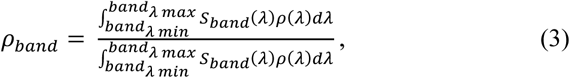

where ρ_band_ is the simulated reflectance of a Sentinel-2 sensor band, *S*_band_(λ) is the sensor spectral response function for the band, and *band*_λmin_and *band*_λmin_ are the minimum and maximum wavelengths where *S*_λmin_(λ) is greater than zero, and ρ(λ) is the simulated spectral information for a given wavelength λ. Examples of the LUT spectra transformed to Sentinel-2 specifications with corresponding FMC values are shown in Fig. S1. In the RTM inversion, we included the Normalised Difference Infrared Index (NDII), calculated from bands VNIR4-842nm and SWIR2-2190nm, alongside reflectance bands, as NDII is sensitive to vegetation water content and has been shown to improve estimation accuracy [58].

The inversion itself was based on the spectral angle, merit function which finds similar spectra by minimising the differences between observations and simulations, following equation 6:

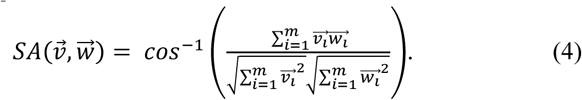

where the reflectance spectrum of each pixel (*W*^→^) was compared in multidimensional spectral space with each of the LUT spectra (*v^→^*, no. of dimensions = *m* = no. bands and indices). The function is calculated in radians, is insensitive to albedo or illumination features in the satellite data [59], and is sensitive to reflectance changes with plant dehydration [35]. To calculate the final prediction, we used the FMC value from a set of simulations the most like each observation (i.e. solutions).

To assess the performance of the RTM inversion, the standard deviation of the set of optimal FMC solutions was calculated as a measure of the retrieval uncertainty and compared with the FMC predictions, observations and residuals [35]. Similarly, we calculated the uncertainty in LAI retrieved from the LUT to assess potential differences in LAI of the LUT to that observed in the field, by taking the standard deviation of LAI of the optimal solutions (according to FMC accuracy and errors).

To minimise unrealistic spectra in the LUT, we tested the effect of applying an ecological filter based on observed FMC and leaf area index (LAI, see Fig. 4), following [35] (i.e. model version M3, which also included the optimisation of solutions as described above). To explore how canopy density might affect reflectance data and RTM inversion, we ran regressions between observed and predicted values of FMC and LAI. We also tested the use of 10 m resolution reflectance data, in addition to 20 m, because this is the native resolution of some Sentinel-2 spectral bands; the remaining bands were interpolated to this resolution during data provision by the custodians [54], [55].

### D. Random Forest Emulation of Radiative Transfer Inversion

The random forest model was trained on a dataset of FMC values derived from the RTM inversion, covering a wide range of forests and woodland types over the period 2015-2024, along with associated Sentinel-2 reflectance bands (n = 7,499, 25% held back for testing). Pixels were randomly sampled from a grid of 20 m resolution, limited to areas with at least 35% foliage projective cover [60], below which accuracy of the RTM inversion was found to be unsatisfactory. We optimised the RF model by tuning the number and depth of trees (n=10 to n=50 and n=5 to n=50) and by conducting a feature importance analysis across different combinations of reflectance bands and indices (Table S1). The indices included NDII which captures foliar water absorption in the infrared, and the normalised difference vegetation index (NDVI), which reflects foliar water via the red edge feature of productive, hydrated vegetation [35]. The RF modelling was implemented in Python using the Scikit-learn, Pandas, Xarray, and Sklearn-xarray packages [61], [62], [63], [64], [65].

### E. Validation of FMC Predictions

For field data pooled across sites, we considered the best FMC prediction from the RTM inversion to be that with the highest explanatory power (r^2^) and low prediction errors in a regression of predicted against observed values, combined with relatively low retrieval uncertainty (i.e. the FMC standard deviation of LUT solutions). High retrieval uncertainty indicates that FMC of solutions drawn from the LUT are not very similar, and this becomes more likely with increasing number of solutions drawn. We calculated Root Mean Square Error (RMSE) between predictions and field observations, including systematic and unsystematic errors therein, to differentiate modelled from uncontrolled contributions to the error. In relatively accurate models, the unsystematic or random errors will be closer to the RMSE and the systematic errors will approach zero. Additionally, we calculated the less ambiguous Mean Absolute Error (MAE) between model predictions and field observations [66]. These error metrics were calculated as follows equation 7:

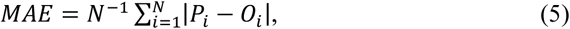

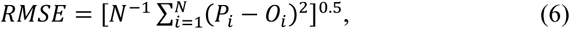

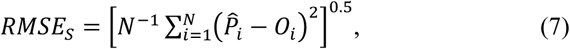

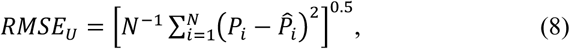

where *N* is the number of observations, *P*_i_is the predicted variable (i.e. modelled FMC), *O*_i_ is the observed variable (i.e. field observed FMC), *P*_i_ is the least-squares regression of *P*_i_ and *O*_i_ These same measures of accuracy and error were used for assessing the FMC predictions of all versions of both the RTM inversion and RF emulator models.

For the RF emulation modelling, we assessed the influence and contribution of predictor bands and indices by analysing the feature importance from candidate RF regressions. This included analysis of the Gini and permutation feature importance, which both involve monitoring the performance of the fitted model as a focal predictor is adjusted across a range of values [62]. The Gini metric alone can be misleading if there are many unique values in the predictors [67]. To identify the best models amongst candidates with varying RF tree structure, we selected the model that maximised the accuracy (r^2^), minimised RMSE and MAE between the RF training data (i.e. RTM inversion) and the RF predictions, while maintaining a relatively fast computation time. To visually assess the predictions, we produced maps of mean, standard deviation and 5^th^ percentile FMC across the complete time series.

## III. Results

### A. FMC of Forests and Woodlands in NSW

The FMC data cube produced using the RF emulator resulted in 2821 unique dates of satellite sampling (07-2015 to 06-2024), over NSW which comprises 61,775 pixels by 51,045 pixels at 20 m ground resolution (i.e. x by y). As an illustrative example of forest and woodland FMC, the year 2019 of the data cube (see Data and Code Availability) was selected due to exceptionally dry climatic conditions and averaged across time. This revealed FMC to be greater near the eastern coast and at higher elevations elevation (e.g. > 600 to 700 m.a.s.l., Fig. 2.)

FMC field observations ranged from 70% to 157 % dry weight, with a mean of 106.1% dry weight (Fig. 3, Fig. S3). This shows that FMC can vary widely in *Eucalyptus* forests and woodlands, and that on average, foliage consists of approximately equal parts water and dry matter. The value of FMC observations generally increased from the climatically more arid sites in the interior of NSW towards the more mesic coastal sites in the east. However, there was one positive outlier in the west that defied the trend because it is a riparian site where the *Eucalyptus camaldulensis* (river redgum) trees sampled had persistent access to ground water. The most abundant species among sites were *E. largiflorens* (n = 5), *E. macrocarpa* (n = 4), *E. albens* (n=3) (TABLE S**3**).

**Fig. 3.**
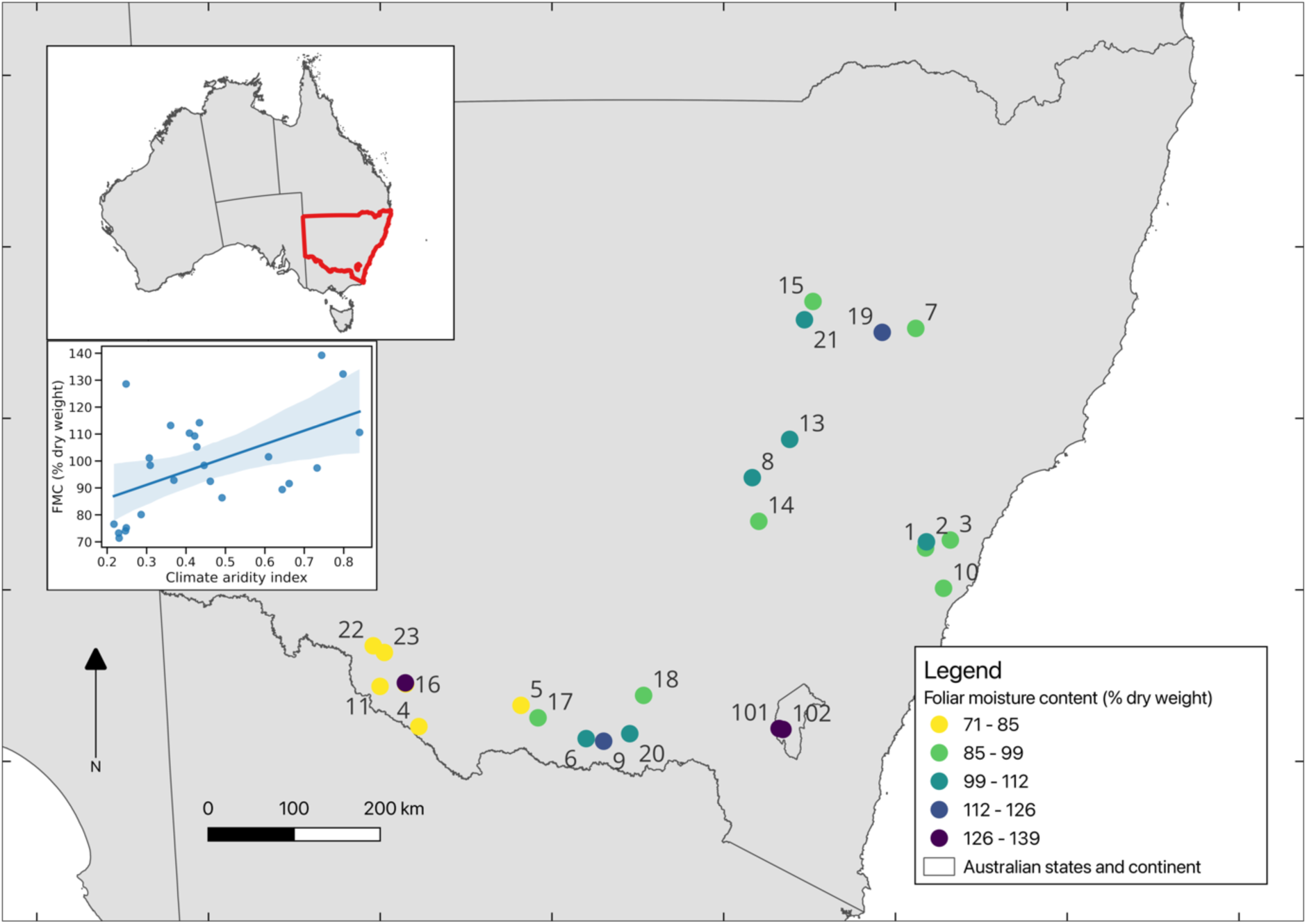
Site foliar moisture content in NSW and the ACT. FMC values may be the mean of multiple observations, if taken, at each site during field campaign in 2022. Site IDs correspond to Table S4 except 101 and 102 which are sites from Yebra et al. [79]. Inset map shows NSW (red highlight) in the whole continent of Australia. Inset regression plot shows the relationship between climate aridity (mean annual precipitation / mean annual potential evaporation) and FMC (% dry weight), with a shaded band indicating the 95% confidence interval.

### B. FMC Prediction Accuracy and Errors

On average, the satellite reflectance measurements were acquired within 2.7 days of the field sample date; for 50 out of 94 field observations there were satellite reflectance measurements available within the required 10-day time window around each field observation date. This loss of field observations was due to cloud cover or atmospheric interference with the reflectance signal of the vegetation during some field sampling days. Foliar moisture content of the tree canopy at field sites was predicted with moderately-high accuracy; the model explained a majority of observed variation (r^2^=0.62, *P<*0.01) and prediction errors were relatively small (RMSE=17.94%, Fig. 4, TABLE 1). This error value is equivalent to 14.39 % on a fresh weight basis. This result demonstrates that FMC can be predicted with small errors at a spatial resolution matching the scale of individual canopies of large trees. (i.e. 20m).

**Fig. 4.**
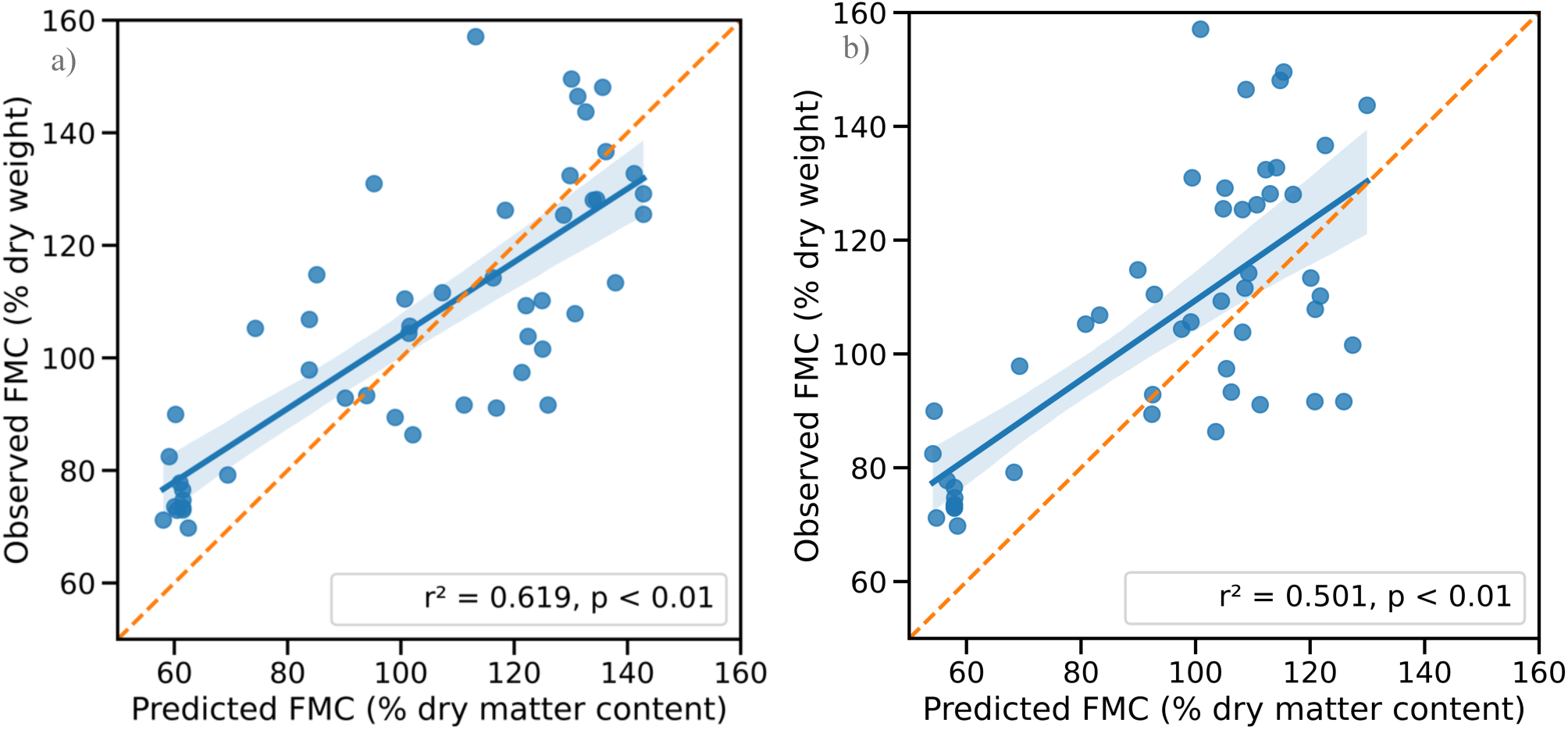
Relationships between observed (FMC_obs_) and predicted (FMC_pred_) foliar moisture content for the two best models: (a) 20 m spatial resolution and (b) 10 m resolution. Orange, dashed line is the 1:1. Note ‘dry weight’ and ‘dry matter content’ are equivalent (Equation 1). Blue solid line indicates the linear regression with blue shading indicating the 95% confidence interval. Each data point represents a single timepoint at each site (mean of 3 leaf samples).

**TABLE 1.** Evaluation of different inversion techniques for FMC estimation based on Sentinel satellite data (versions described in methods section 2.3). n: number of sites in validation of modelled fmc, fmc_sd_: standard deviation of modelled fmc, noptim: optimal number lut spectra to consider in merit function, a: intercept of fitted validation least-square regression, b: slope of fitted validation least-square regression, mae: mean absolute error (%), rmse: root mean squared error (% dry matter content), rmses: Systematic rmse (% dry matter content), rmseu: unsystematic rmse (% dry matter content).

| Model version | N | $FMC_{min}$ | $FMC_{mean}$ | $FMC_{max}$ | $FMC_{SD}$ | $n_{optim}$ | a | b | $r^2$ | MAE | RMSE | $RMSE_s$ | $RMSE_u$ |
| --- | --- | --- | --- | --- | --- | --- | --- | --- | --- | --- | --- | --- | --- |
| M3 | 50 | 52.60 | 99.07 | 145.45 | 30.48 | 57 | -7.2058 | 1.0017 | 0.62 | 15.02 | 17.94 | 7.03 | 18.64 |
| M2 | 50 | 56.69 | 97.29 | 134.21 | 23.80 | 150 | 17.3098 | 0.7539 | 0.50 | 15.42 | 18.64 | 10.56 | 15.36 |
| M1 | 50 | 46.41 | 91.02 | 131.09 | 24.42 | 40 | 11.6504 | 0.7480 | 0.49 | 19.85 | 23.09 | 16.22 | 16.44 |

There was some underprediction of FMC at the lowest observed values. The RMSE had a ratio of systematic to unsystematic error of 7.0:18.7 (TABLE 1). This means that the unsystematic (i.e. random) error is much larger than the systematic (i.e. controlled) errors and is approaching the overall RMSE, a characteristic of good models [68], while the systematic error is closer to 0, as is ideal.

When comparing the predictions of FMC from RTM models using reflectance data at different ground resolutions, the 10 m resolution model explained less variation in FMC_obs_ than the 20 m resolution model (r^2^ = 0.50 compared to r^2^ = 0.62, Fig. 4Fig. 4b). The finer spatial resolution model had a similar optimal number of solutions (n = 37) and slightly higher errors (RMSE = 21.81, and MAE 18.95), compared to the 20 m version of the model. The cluster of underpredicted values at the lowest FMC_obs_ was present in both models, suggesting that this error component was not related to the spatial resolution of reflectance data. For the 20 m spatial resolution model, the range of predicted FMC was 53% to 145%, with a mean of 99%. This model was found to generally underpredict FMC, particularly in a group of observations with low FMC_obs_ (∼ - 20%, Fig. 4)

### C. LUT Ecological Filtering

There was a significant positive relationship between observed LAI and FMC (r^2^=0.22, *p =* 0.02, Fig. 5Fig. 5a). Filtering the LAI-FMC space of the LUT with the observed relationship (±10%, Fig. 5Fig. 5b), reduced the LUT from *n* = 4482 to *n* = 3015 (Fig. 5Fig. 5c and Fig. 5d, respectively). Therefore, ∼1/3 of LUT entries had unrealistic LAI-FMC relationships, mostly those of high LAI and moderate to low FMC (i.e. top-left of Fig. 5c).

**Fig. 5.**
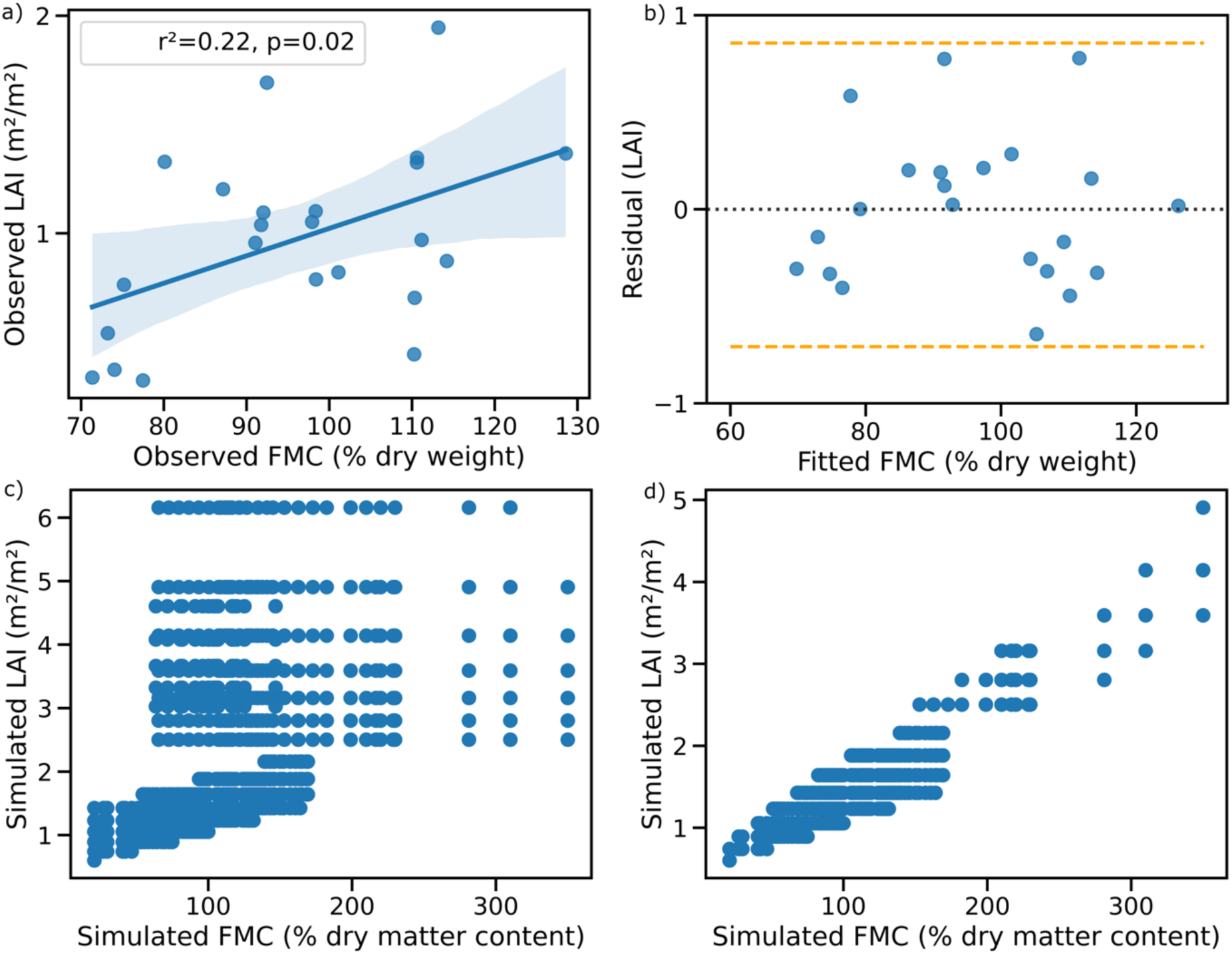
Ecological filtering of the LUT. a) Relationship of observed LAI by observed FMC: y = 0.012493 x - 0.227067, b) residual plot of observed LAI and observed FMC, including the maximum +10% and minimum −10% (dashed, orange lines), c) scatterplot of simulated LAI against FMC (i.e. LUT parameters) b) simulated LAI against FMC after filtering for the ecosystems under study (i.e. by dashed orange lines in b). Note the different x and y axes for plots showing similar variables.

### D. Number of Solutions in the Inversion

The optimal number of solutions in the RTM inversion was *n* = 57 (Fig. S4). Including more solutions reduced the RMSE marginally, however it did not increase the accuracy (i.e. r^2^). An optimal number between approximately *n* = 200 and *n* = 300 solutions also resulted in a good relationship comparing predictions and observations (r^2^=∼0.61, RMSE=∼16%, Fig. S4). These results suggest that there is more than one good set of solutions, however fewer but not singular solutions produce the best prediction of FMC.

### E. Retrieval Uncertainty

The magnitude of the retrieval uncertainty for FMC was found to be significantly negatively related to the observed, and the predicted FMC (Fig. 6Fig. 6b & c). There was no relationship between the uncertainty and the residuals (Fig. 6Fig. 6a), indicating that the error is not being primarily driven by variation in FMC of the LUT. The retrieval uncertainty for FMC increased in a power curve with increasing number of solutions (Fig. S5). This indicates that the RTM inversion is less certain when including more solutions, despite reduced errors with more solutions (Fig. S4).

**Fig. 6.**
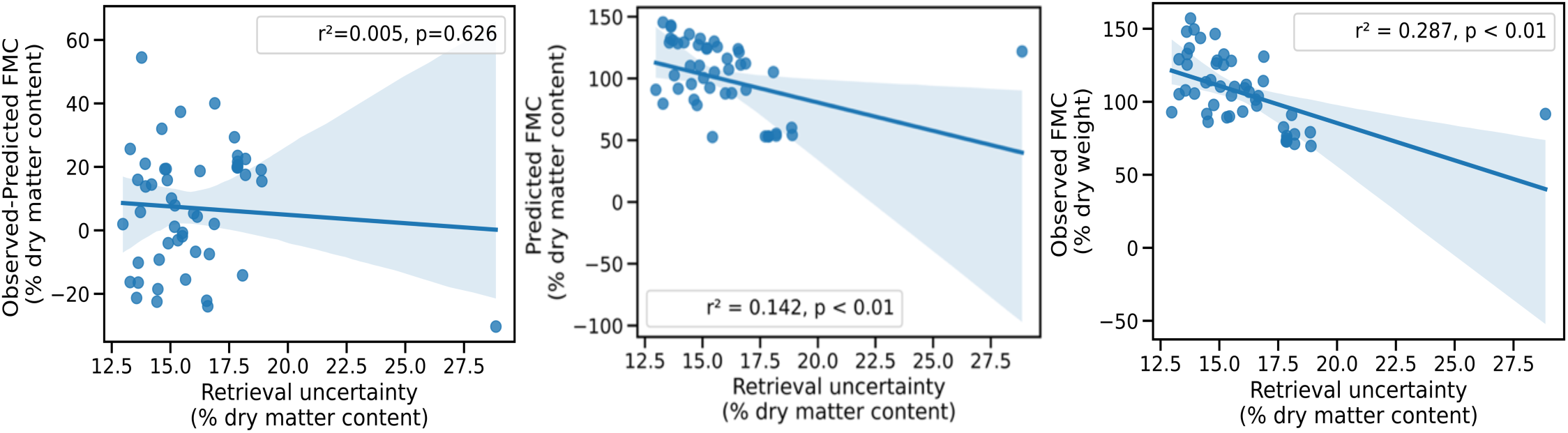
Accuracy of the inversion illustrated by retrieval uncertainty (calculated as standard deviation of the 57 solutions used in the best model) plotted against the residuals from the validation (a), also against field measured (b) and predicted (c) FMC. Note y-axes are different amongst plots.

### F. Canopy Structure

There was a weak positive relationship between FMC_obs_ and LAI_obs_ (Fig. 7Fig. 7a), that is FMC tends to be lower at low LAI sites. There was a stronger positive relationship between LAI_obs_ and predicted FMC (Fig. 7Fig. 7b). These results suggest that reflectance-based measurements depend on the amount of leaf area contributing to the signal. They also suggest that the LUT used for predictions may be biased—overemphasizing the influence of LAI on predicted FMC.

**Fig. 7.**
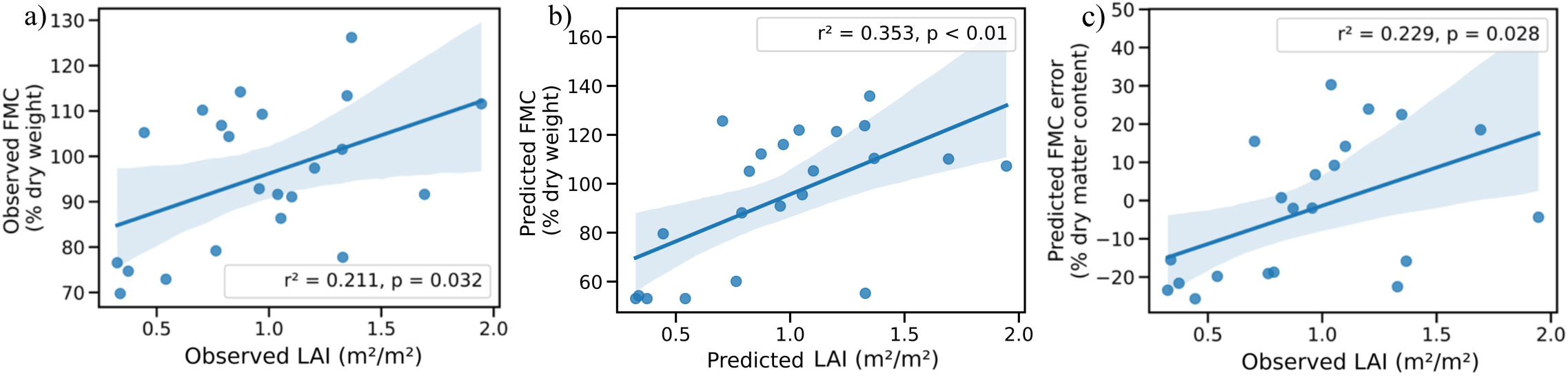
Relationships between a) observed FMC (FMC_obs_) and observed LAI (LAI_obs_), b) predicted FMC (FMC_pred_) and predicted LAI (LAI_pred_) and c) FMC prediction error and observed LAI (LAI_obs_), including regression lines and shaded 95% confidence intervals.

There was a significant positive relationship between the relative prediction error of FMC and LAI (*p=*0.028, Fig. 7c). This suggests that underprediction is associated with sparser canopies and conversely that there may be some overprediction of FMC associated with denser canopies. This result was confirmed by replacing LAI_obs_ with a satellite-based estimate of foliage projective cover [60] that also produced a significant positive relationship between the relative prediction error of FMC and LAI (*P<*0.01, Fig. S6).

Variation among LUT solutions for LAI tended to increase with LAI_pred_ (Fig. 8), alongside a reduction in FMC prediction errors. This can be seen in Fig. 5c and Fig. 5d, by the slope of the lower bound of the FMC-LAI relationship, which was not an effect of filtering the LUT by site FMC-LAI. These results indicate the LUT is restricted at low LAI values with respect to FMC_obs_.

**Fig. 8.**
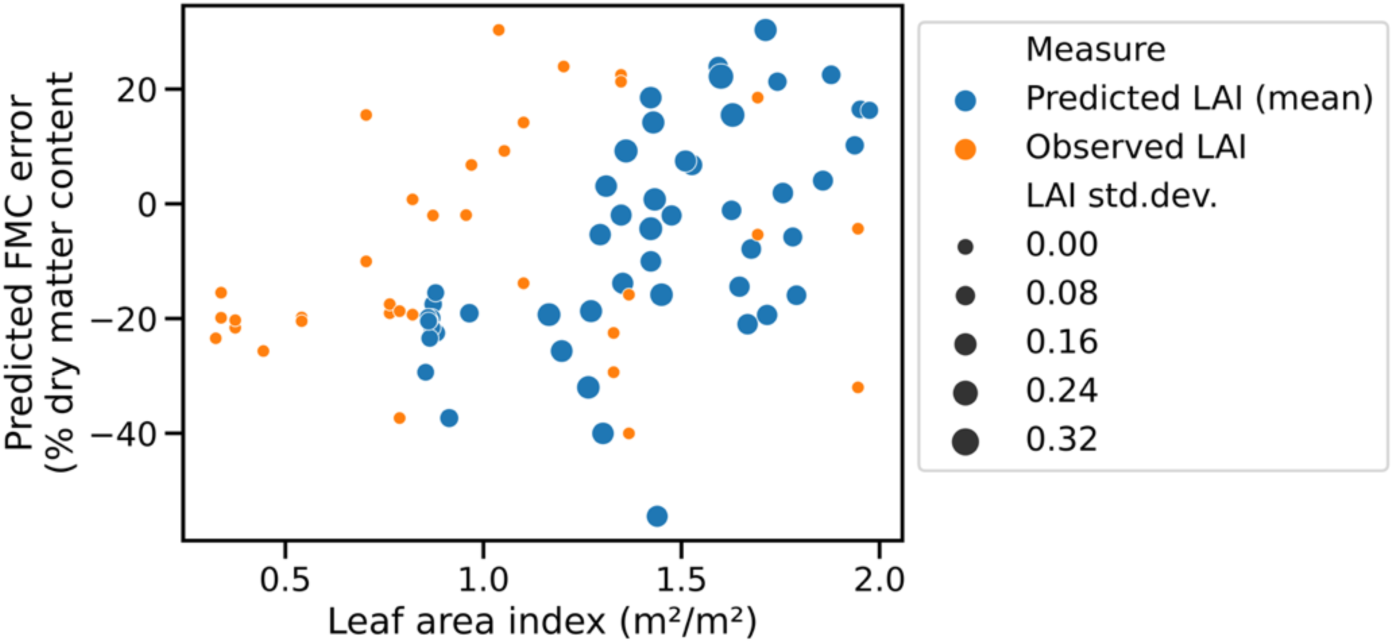
Bubble plot of FMC prediction error against LAI, observed (orange) or predicted (blue, i.e. average for n=57 spectra retrieved from LUT). Bubble size is the standard deviation of LAI of retrieved spectra per observation (i.e. only relevant to predicted LAI (blue).

### G. Random Forest Emulator of RTM

The RF model emulated the RTM predictions of FMC with an error of approximately 10% (RMSE) and a high coefficient of determination (r^2^ = ∼0.9, Fig. 9a). This result demonstrates that substituting a physically-based FMC model utilising satellite reflectance with a machine-learning model trained on that same data source leads to a reduction in the overall explained variance in FMC of 10% (i.e. inverse of r^2^).

**Fig. 9.**
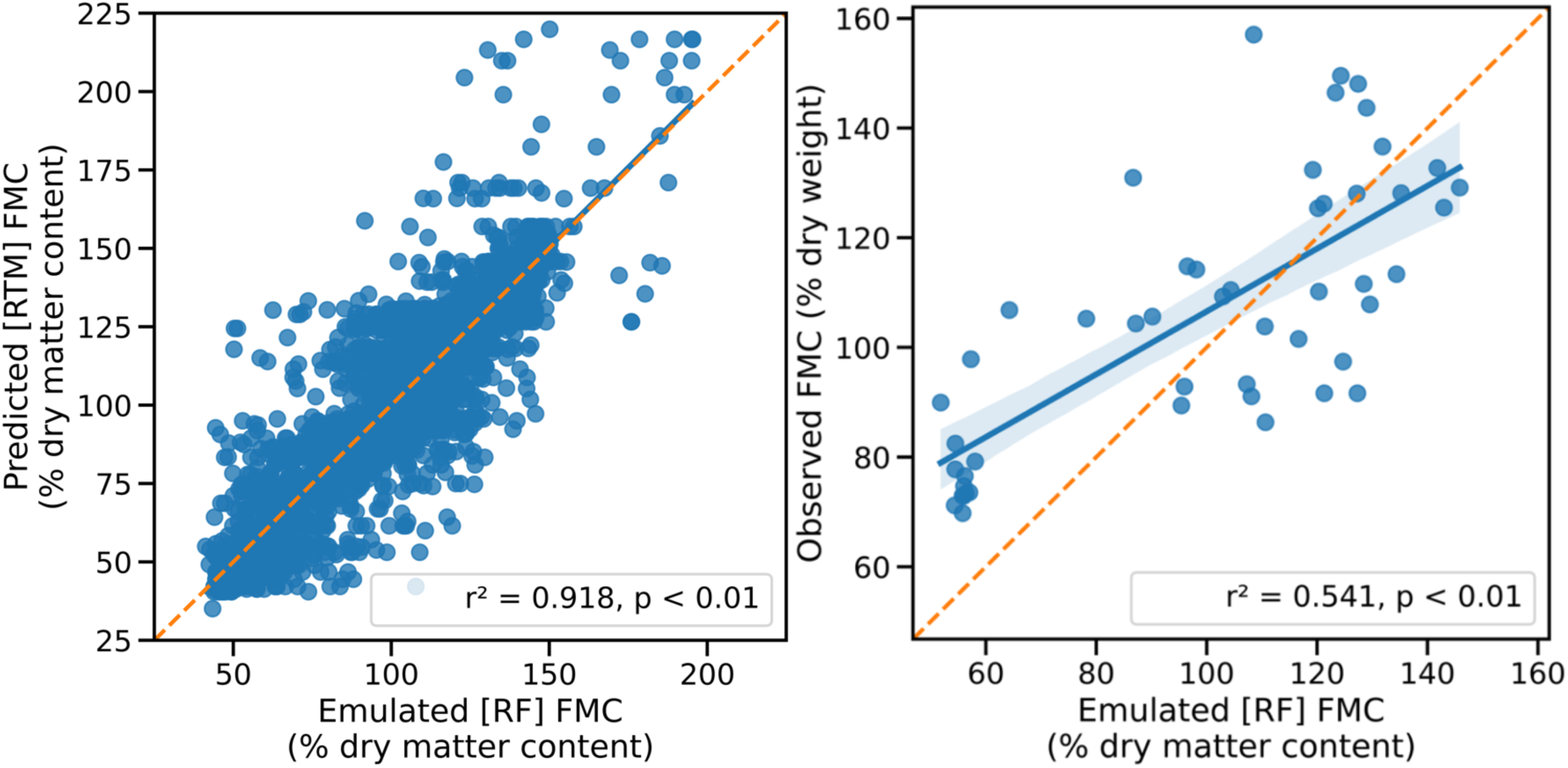
(a) Relationship between FMC predictions from the Radiative Transfer Model (RTM) and Random Forest (RF) model on the testing dataset (n = 1,874). (b) Relationship between RF-predicted FMC values at field sites (n = 50) and corresponding FMC observations. In both panels, the dashed line represents the 1:1 line (line of perfect agreement), while the solid blue line shows the fitted linear regression, with the shaded area indicating the 95% confidence interval.

The emulator model of FMC, when validated against field observations, still had acceptable accuracy (r^2^=0.54) and RMSE of 21.77% (RMSE, Fig. 9b). As expected, the performance of the emulator model is slightly less than the RTM inversion on which the emulator model was built when using the same field data in comparison (i.e. Fig. 4, ^2^=0.62, RMSE=17.94%, *P<*0.01). The underprediction by the emulator model at low field values was consistent with the original model. The computation time for the RF emulator model was generally > 200 times faster than the RTM inversion.

## IV. Discussion

Effective decision-making in conservation and fire management on topics such as assessing habitat quality for arboreal folivores and forest flammability rely on accurate information about the spatiotemporal variation of foliage moisture content (FMC). In this study we have tested the inversion of an RTM to predict FMC using Sentinel-2 reflectance data and produced a high-resolution data cube of forest FMC for NSW, Australia. By accounting for canopy structure, using field measured LAI and an FPC product, we were able to improve FMC predictions relative to previous similar models. The results show improved accuracy in predicting forest and woodland FMC with higher explanatory power (r^2^=0.62) and lower error (RMSE=17.94%), compared to previous studies: Jurdao et al. [40] reported r^2^=0.5 and RMSE=30% for mixed woodlands in Spain, while Yebra et al. [35], found r^2^=0.17 and RMSE=32% for forests across Australia. Notably, our study achieved these improvements despite using the same LUT developed from mixed woodlands in Spain as in both previous studies. The improvement may mostly be driven by the higher ground resolution of reflectance data used, particularly in the water-sensitive SWIR bands, which approaches the scale of individual tree canopies.

Previous work in Australian forests that used the same LUT as in the present study showed similar errors across observed FMC values, with higher error at low FMC [33]. This similarity occurred despite the use of different reflectance data sources ([33] used MODIS). Therefore, these errors may arise from the parameterisation of the original RTM and resultant LUT, which were based on mixed woodlands in Spain [40]. Limited representativeness of RTM inputs has been recognised previously; Ma et al. [69] dealt with the issue by using global databases of plant traits and soil reflectance to parameterise the PROSAIL model before inversion with Sentinel-2 and other reflectance data.

We found retrieval uncertainty, which is a measure of the deviation of a parameter within the set of retrieved LUT solutions [35], and FMC_obs_ had a negative relationship so that uncertainty was higher at lower FMC. The relationship is the opposite of that found in Yebra et al. [35], where the RTM was inverted on MODIS data to predict FMC in forests as well as grass and shrublands. Further, we found a stronger relationship overall of FMC retrieval uncertainty with FMC_pred_, as opposed to FMC_obs_, that suggests the retrieval uncertainty is being driven by the model inversion rather than variation in the reflectance data. The higher uncertainty at lower FMC values should be noted if the output data cube is to be used in studies where lower values are likely to be of interest (e.g. tree dieback or fire fuels). At our field sites, the LAI of the eucalypt-dominated ecosystems was important for both the reflectance signal and the RTM inversion. The stronger relationship of LAI_obs_ with FMC_pred_, rather than that with FMC_obs_ suggests that our model reinforces the influences of LAI influence on FMC. This might be the source of the retrieval uncertainty coming from the inversion (see above), which would have contributed increased RMSE. Such reinforcement may be due to a lack of solutions with relatively low LAI coupled with moderate to high FMC (i.e. ³ ∼150%, Fig. 5c, d). For instance, the interquartile range of LAI in the LUT, before filtering was 1.23 to 1.64 m^2^/m^2^, whereas at our sites it was 0.7 to 1.3 m^2^/m^2^. This difference occurred despite the LUT having been produced in such a way that lower LAI was more comprehensively sampled [40], though the LUT was filtered to the LAI-FMC domain encompassing our field data. This LAI-FMC mismatch might also be explained by the Mediterranean, species used to produce the LUT [40], not holding as high moisture values with low LAI as the temperate and semi-arid *Eucalyptus* species studied here (i.e. less sclerophyllous). Some of our sites had LAI < 0.4, while minimum LAI of species used in producing the LUT was 0.6, though the latter was taken from a few sources of differing methodologies [70]. However, trees have limits in their spectrum of relations between leaf water content and leaf area so that chances of higher water content with lower leaf area, are generally low [71].

Improvements to FMC estimation in south-east Australia could be made by undertaking new forward radiative-transfer modelling. Despite the LUT containing only one *Eucalyptus* species of nine LUT species total, the FMC predictions were more accurate than in previous studies based on the same LUT. However, developing a new forward stage of RTM modelling—such as a LUT based exclusively on species from our study region with a more representative FMC–LAI relationship—could further improve estimation accuracy. Other improvements could include fixing LAI in the RTM inversion with a dedicated, remotely-sensed LAI product, or equivalent measure [35], [40], to reduce the issue with representativeness of LAI-FMC in the LUT, described earlier. However, the LUT may first need to capture the full LAI-FMC domain of interest, else this method would provide limited improvements in FMC predictions. However, caution must be taken as LAI estimates from satellite data potentially propagate modelling errors into predicted FMC [35].

Another explanation for the underprediction of lower FMC, aside from the LAI-FMC relations, may be a reduced canopy signal at low LAI. In savanna woodlands, Fuller et al. [72] found that the tree canopy contribution to reflectance, in terms of the NDVI, decreased below ∼60% tree cover. Of our 23 sites, five had tree cover greater than 60%. The sites with the lowest tree cover (e.g. Yanga NP, Mallan, Moulamein, Mt. Arthur and Yanga SCA) tended to underpredict FMC by ∼15 % DMC, averaged across sites. However, underprediction of FMC also occurred at some sites with tree cover exceeding 60%. It is possible that a lack of canopy signal combined with the unrepresentative LAI-FMC relationship in the LUT influenced the underprediction of low FMC in our model, and that used by Griebel et al. [33] which share the LUT.

Despite issues of canopy density, more accurate FMC predictions were achieved partly because of higher resolution reflectance data, relative to previous similar studies (e.g. 500 m resolution, [35], [40]). Since these previous studies share the same LUT, we can assess the effects of the different spatial resolutions on the accuracy of FMC prediction. The spectral signal of 20 m Sentinel-2 pixels will likely only capture a few individual trees and include few tree species resulting in a less mixed spectral signal [73]; depending on habitat type, eucalypts typically have crown diameters of less than 20 m but they may be even larger [74]. Furthermore, gaps exist within and between tree canopies, through which understorey reflectance contributes to the signal received by a satellite sensor [75]. However, this contribution is less likely with finer spatial resolution data because gaps can be isolated from the target in separate pixels and then masked out, as was done here with the foliage projective cover data. The apparent benefit of increased spatial resolution was not seen however in our comparison of two resolutions of Sentinel-2 reflectance data (Fig. 4Fig. 4), where the model based on 10 m resolution reflectance data showed no improvement over the model based on 20 m resolution data. This may be due to the Sentinel-2 sensor limits of 20 m native resolution in the two SWIR and one VNIR bands, which were interpolated to 10 m [54]. Only visible and some NIR bands have 10 m native resolution. SWIR bands are relatively important in spectral response to FMC [47] and so the lack of actual increase in resolution in these bands will show little improvement in the spectral response to FMC. Additionally, increasing resolution to 10 m effectively reduces the representativeness of the field measurements by potentially excluding sampled trees that occurred in our 20 m x 20 m field plots.

The size of the forest area covered by field sampling plots affects the degree of spatial representation of a tree attribute. FMC may vary within single trees and amongst the individuals within the plot. For instance, Griebel et al. [33] found that tree average FMC within 25 m radius plots could vary by as much as 59% dry weight; they also showed within-tree variation in FMC of as much as 55% dry weight (91 to 146% dry weight). In the present study, samples from 20 m x 20 m plots were made up of three subsamples, usually from different trees, with the deviation of FMC between these samples being up to 26% dry weight at some sites. This within-plot variation is only marginally larger than the error of the FMC model overall. Increasing the sampling intensity to more than three samples may reduce this difference. However, we can only assess error relative to our field measurements, if there is sampling error associated with field measurements, then RTM predictions might be closer to reality, because they are capturing reflectance off the entire canopy in each pixel. Furthermore, we did use greater field sampling intensity than in previous studies which had larger plot sizes [35], [76] and few samples can represent an aggregated plot value [77].

Our model based on Sentinel-2 imagery included more information in the NIR and the same number of bands in the SWIR portion of the spectrum compared to previous models based on MODIS data [35] (see Table S1 for band comparisons). Variation in SWIR wavelengths have the strongest response to FMC, however there is also some response to FMC in NIR wavelengths. The SWIR bands tend to increase in reflectance with FMC due to variation in vegetation dry matter content, with its contribution to reflectance variability being expressed at ∼750 nm to ∼1300 nm [78]. The MODIS sensor has only one band in the SWIR region compared to four in the Sentinel-2 sensor used in our model (Band 1 and Bands 7, 8, 8a, 9 respectively, Table S1). This increase in spectral resolution, alongside the increase in spatial resolution discussed earlier explains some of the improvement in results from previous models.

Our new FMC data cube provides unprecedented accuracy, resolution and coverage of east Australian forests and woodlands, opening new opportunities for ecological applications. Desiccation of vegetation i.e. declines in FMC, via the disruption of tree hydraulic function implies a loss of water from the ecosystem as seen in widespread instances of canopy browning or dieback [3], [7], [31]. This process may limit the capacity of arboreal folivores to thermoregulate and increases the flammability of forest fuels [13], [17], [27]. Overall, the FMC error of 18% observed here in forests and woodlands is relatively small, compared to the range of variation observed in FMC_obs_ of 70% to 160% (Fig. S3, i.e. converted to WC, error = 17% and range = 41% to 62%). The error is also relatively small compared to the observed range of forest fuel dryness (∼63%-140%) and the distinct FMC thresholds (99%-106%) associated with increased levels of fire activity in eucalypt forests of SE Australia [8]. Nolan et al. [8] also noted that FMC estimates aggregated over large pixels, inevitably miss within-plot variation in live fuel moisture content due to variation in terrain and vegetation types. The FMC predictions produced here at 20 m resolution are more likely to capture fine-scale variation in FMC that may facilitate, or hinder fire spread. The temporal variation in FMC of Australian forests and woodlands has been shown to be from 75% to 223% dry weight [23], [24] and 88% to 189% dry weight [79]. Our model, with an error of ∼18%

DMC, should therefore be able to capture such fluctuations in FMC. In animal ecology, FMC may become a limiting factor for folivores during intense or prolonged droughts and heatwaves. Dietary water is usually sufficient for hydration and thermoregulation under most climatic conditions (e.g. for koalas, [80], [81]). The value of this limit in koala forage may be ∼120% dry weight but it is not well established [25]. We expect that FMC declines exceeding 20% dry weight, i.e. the model error, would be required in most places to approach this limit. Therefore, locations and periods in which this limit is approached can be identified through the FMC modelling presented in this study.

Moreover, clear temporal and fine-scale spatial trends are apparent in the FMC data cube modelled here and will be analysed and discussed in more detail elsewhere. These data offer a very useful, relative measure allowing the identification of persistent landscape gradients of vegetation moisture and thus allowing for investigation of the underlying drivers of these patterns. The data facilitate the remote sensing of ecological drought (e.g. [82]), of conditions preceding canopy browning and tree death (e.g. [83]), and of critical moisture levels in live forest fuels (e.g. [35]. It may also allow further testing of theories of leaf economics in which leaf water is an important trait [48]. Critically, it can also allow the identification of locations in the landscape which maintain locally high FMC even in periods of local water stress. These are sites of high and increasing conservation value as climate refugia for many animals in the context of drought, heatwaves and possibly fire. Identifying, protecting and ensuring connectivity to this refugia should be considered a management priority for the conservation of forest-living fauna and flora.

## V. Conclusion

This study showed an improvement in the prediction accuracy of foliage moisture content in woodlands and forests of SE Australia. This was achieved using Sentinel-2 satellite data in an inverse radiative transfer model with forward modelling taken from existing work, then ecologically filtered. The satellite data used was of finer spatial resolution than in similar previous modelling. Our analysis showed a general improvement in FMC estimation compared to previous methods, though there is also a tendency to underpredict at very low FMC. We also presented a data cube of the final FMC model applied across NSW, Australia. The high-resolution FMC data cube provides a valuable resource for further studies in animal, fire and plant ecology, including of climatic effects on animals and landscape effects on forests.

## Data and Code Availability

The FMC maps and time series data cube are available as tiled GeoTiff (maps only) and NetCDF files (maps and data cube) from the authors’ institutional repository at https://hie-pub.westernsydney.edu.au/projects/fmc_modelling/. The projected coordinate system of the tiles of data is NSW Lambert Conic Conformal (2SP, i.e. EPSG:3308). Each tile covers a 5000 x 5000 20 m pixel area (100 x 100km) for a total of 102 tiles covering the Australian state of NSW as shown in Fig. S7, (includes two partial tiles). Each tile is named by the coordinate of the north-western (top-left) corner of each area. Each native tree cover pixel in the data cube includes samples between early 2015, when the Sentinel-2 mission began, and the present (mid-2024, *n*=∼400 to *n=*∼1000 timepoints), with higher numbers in the overlaps between satellite tracks and in regions with fewer cloudy days.

It is important to keep in mind that the FMC data cube presented here incorporates reflectance data collected during large disturbances (e.g. forest clearance, fire). As such, high variation (Fig. S8), for example in the Blue Mountains near Sydney, may reflect not only variation in the intact canopy but of the loss and regrowth of the canopy itself during and after fires, including potentially an understorey signal (e.g. 2019/20 fires, [7]).

When using the full timeseries of FMC data across different ecosystems, a standard score (i.e. z-score) or a ranked percentile, should be considered. Such a measure will highlight changes in FMC relative to the mean for each pixel, reducing the effect of spatial differences in absolute FMC estimates [84]. Code for the RTM, emulator modelling and production of the FMC data cube can be found at the following webpage: https://github.com/ikotzur/fmc_modelling_nsw. The field data produced for this study is also available as raw observations in the GLOBE-FMC dataset version two [36], which is a set of FMC measurements primarily collected for fire ecology (i.e. fuel or live fuel, rather than foliar, moisture content).

---

**Ivan Kotzur** received the B.Sc. (Hons) degree in environmental remote sensing from Australian National University, Canberra, Australia. He has submitted a thesis for the Ph.D. degree in ecological remote sensing and animal ecology with the Hawkesbury Institute for the Environment, Richmond, Australia. His research interests include forests, eucalypts, koalas, fire, ecology and remote sensing.

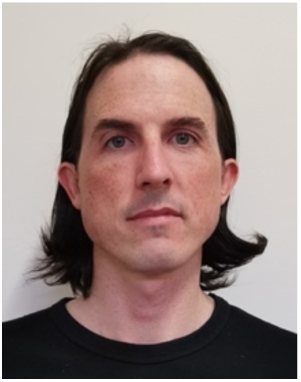

**Professor Matthias Boer** received his Ph.D. in physical geography from Utrecht University in 1999. Since receiving his Ph.D., he has held research positions at the Spanish National Research Council (CSIC), The Commonwealth Scientific and Industrial Research Organisation (CSIRO) in Australia, The University of Western Australia and Western Sydney University; he currently is professor of fire ecology with the Hawkesbury Institute for the Environment. His research work focuses on modelling fire regimes, forest flammability, the effectiveness of fire management, and forecasting of fire risk.

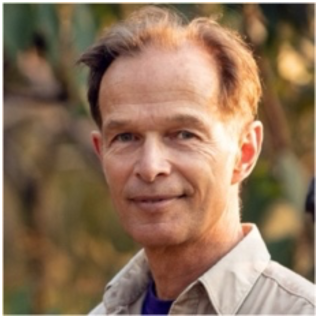

**Professor Marta Yebra** is a Professor in Environmental Engineering at the Fenner School of Environment & Society and the School of Engineering. She is the Director of the Bushfire Research Centre of Excellence supported by ANU which aims to protect Australia from catastrophic bushfires. She is also Mission Specialist at the ANU Institute for Space. Her research focuses on developing applications of remote sensing for the management of fire risk and impact at local, regional and global scales.

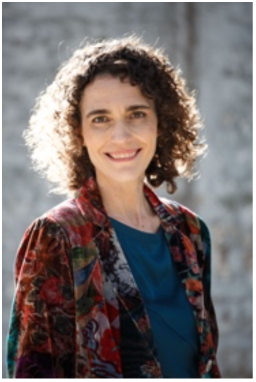

**Associate Professor Ben Moore** is a nutritional and chemical ecologist at the Hawkesbury Institute for the Environment at Western Sydney University. His research focuses on the evolutionary and environmental drivers that shape the nutritional and chemical quality of plants that serve as food for animals, and how animal individuals and populations respond to the resulting nutritional landscapes. In the Australian context, arboreal marsupial folivores like the koala are of particular interest because of their highly evolved and specialised interactions with the eucalypts that dominated forest ecosystems.

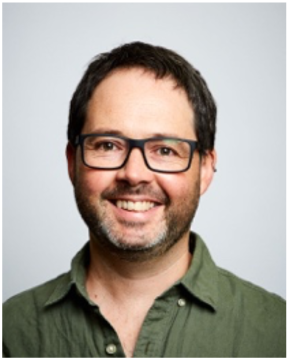

## Supplementary Material

### A. Example Spectra from the LUT

**Fig. S1.**
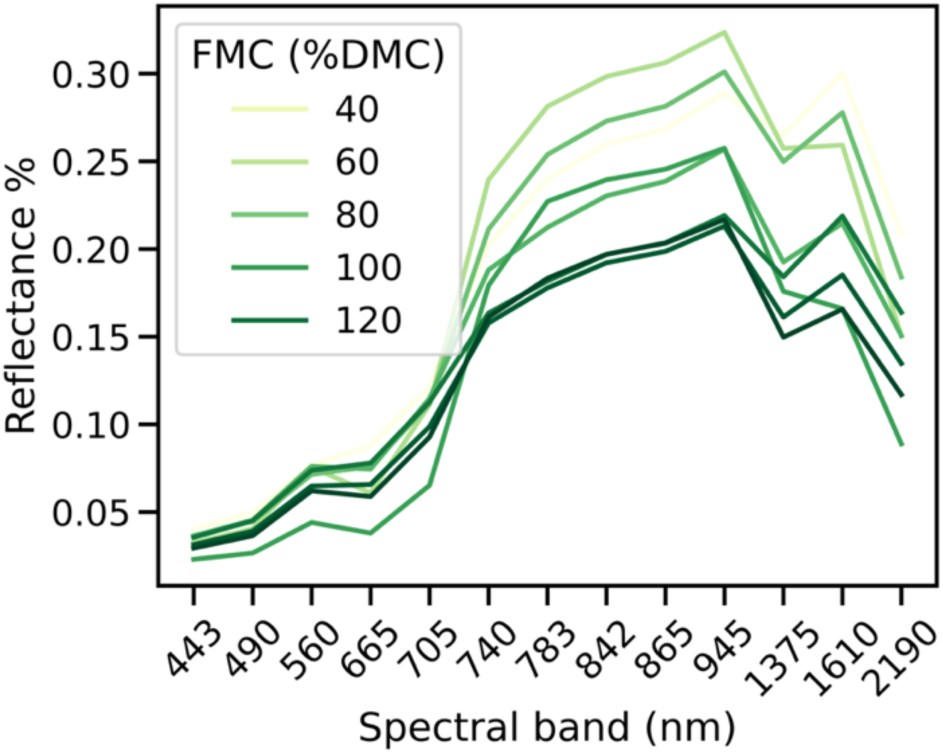
Ten random spectra from the Look-Up Table (LUT, [35]) with corresponding FMC values. Spectra have been convolved to match Sentinel-2 specifications from the full-spectral resolution wavelengths. Total LUT size n = 3015. Note x-axis is non-continuous.

### B. Mean FMC in NSW

The map of mean FMC across the timeseries (Fig. S2) shows that forests and woodlands in NSW tends on average to be higher nearer the coast, in the east, and in the areas of highest elevation. Also, FMC is relatively high along river systems in the west of the state. Central areas inland of the Great Dividing Range tend to have linear patterns of higher moisture content, probably near water courses, whereas closer to the coast, extensive patches of high FMC exist and in the western slopes region these are, apparently, in relatively elevated hill and mountain ranges or mountains.

**Fig. S2.**
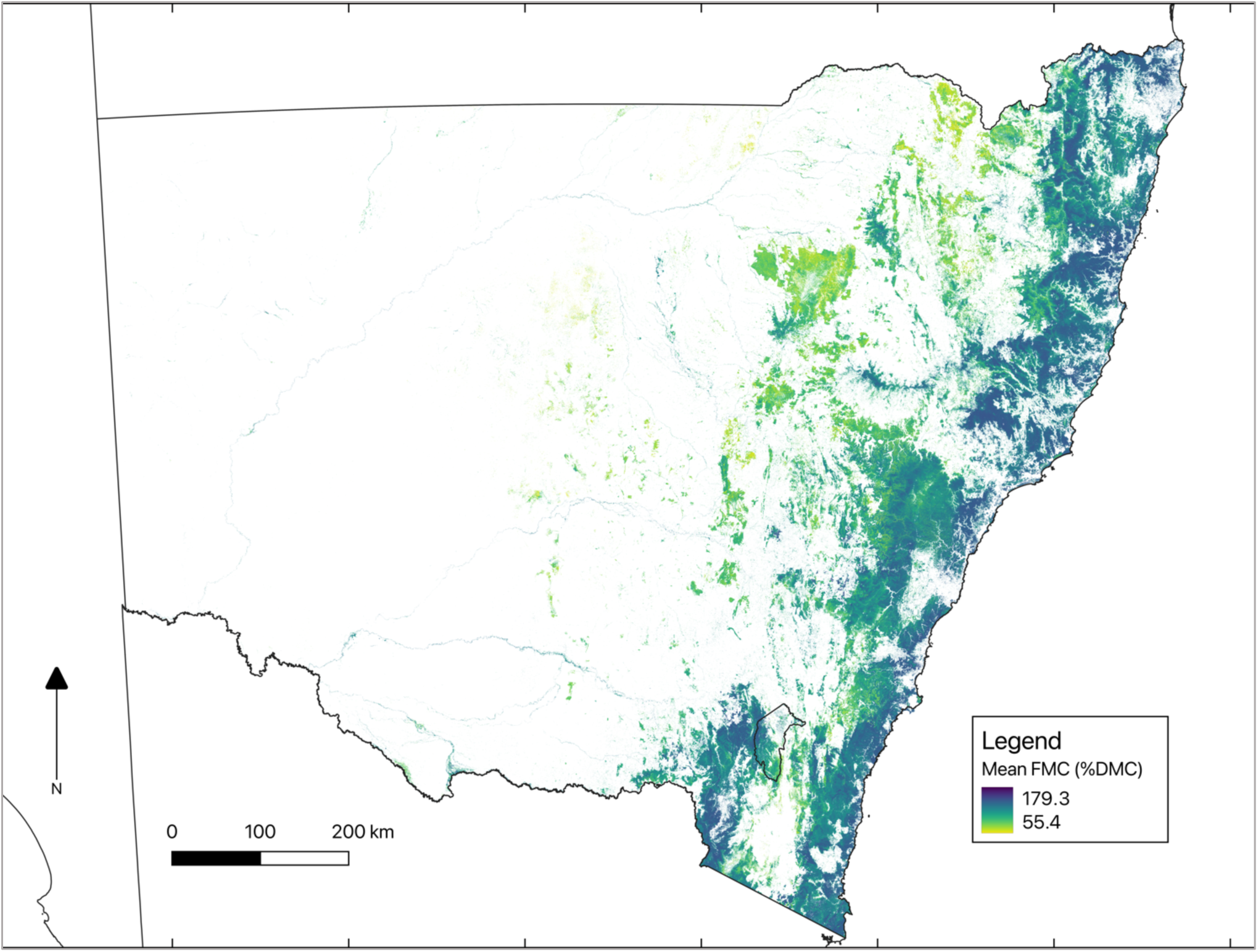
Mean foliar moisture content (FMC, % dry matter content [DMC]) in forests and woodlands of NSW (2015-2024).

### C. Additional Field Data

**Fig. S3.**
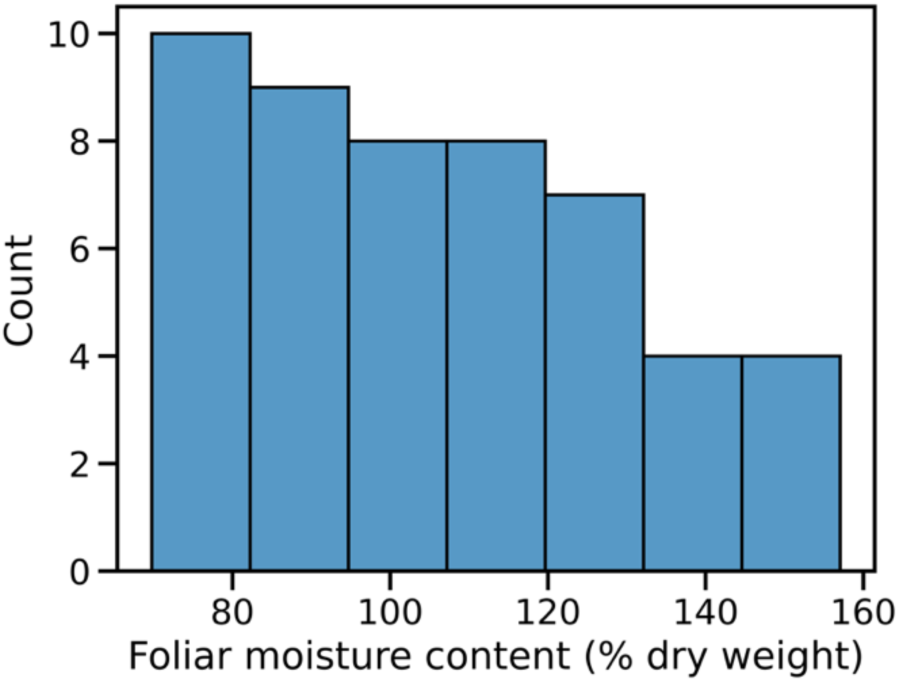
Distribution of foliar moisture content at field sites. n = 50. Mean = 106.1 % dry weight, median = 105.4 % dry weight.

### D. RTM Inversion Evaluation

**Fig. S4.**
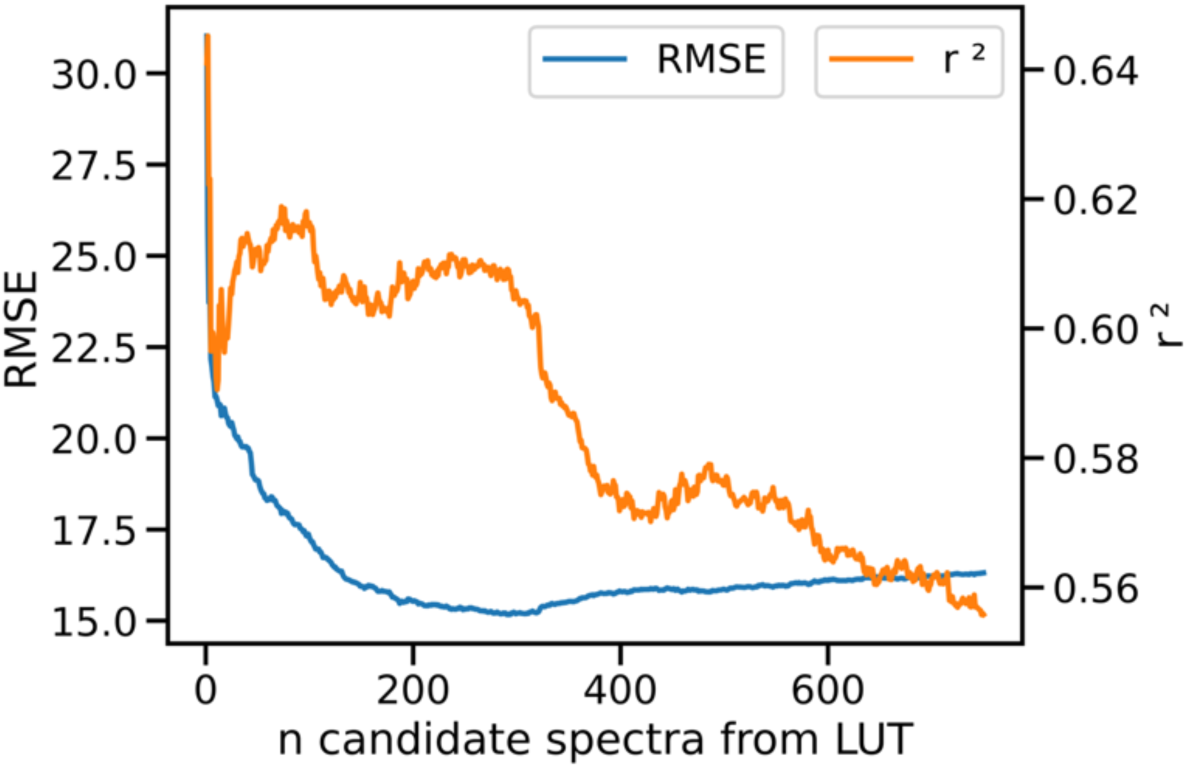
Changes in correlation coefficient and RMSE when altering n most similar spectra from merit function in FMC prediction. RMSE: blue, left y-axis. r^2^: orange, right y-axis.

**Fig. S5.**
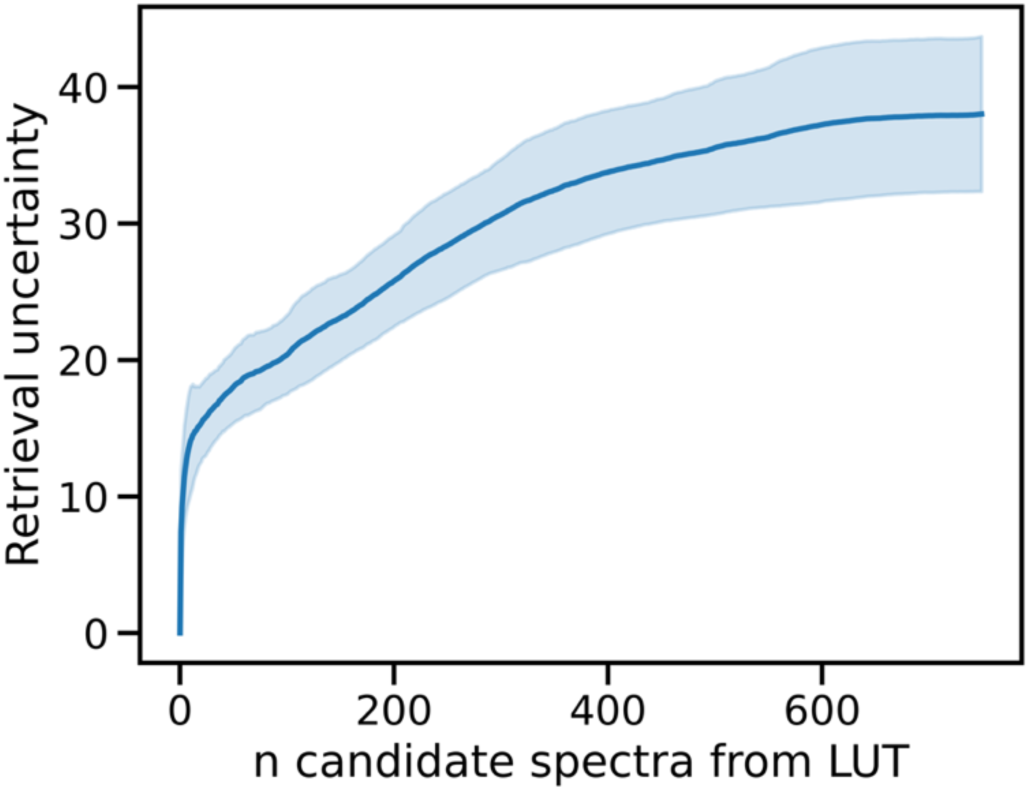
Changes in retrieval uncertainty when altering n most similar spectra from merit function in FMC prediction. Retrieval uncertainty is the standard deviation of FMC of candidate, LUT spectra (i.e. solutions) used in the final FMC prediction (i.e. median of solutions). Error bars are the standard deviation of the retrieval uncertainty (i.e. of all observations).

**Fig. S6.**
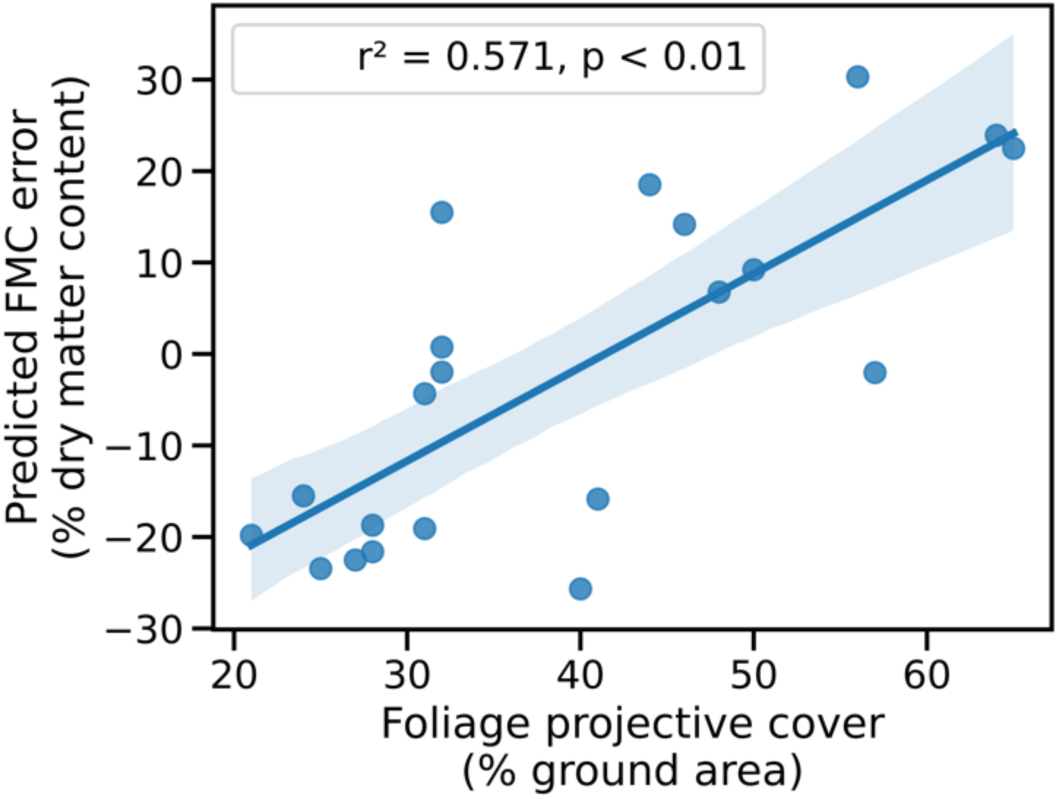
Relationship of errors of FMC prediction with foliage projective cover (FPC), includes linear regression line with shaded 95% confidence interval.

### E. Coordinates of tiles in FMC Dataset across NSW

**Fig. S7.**
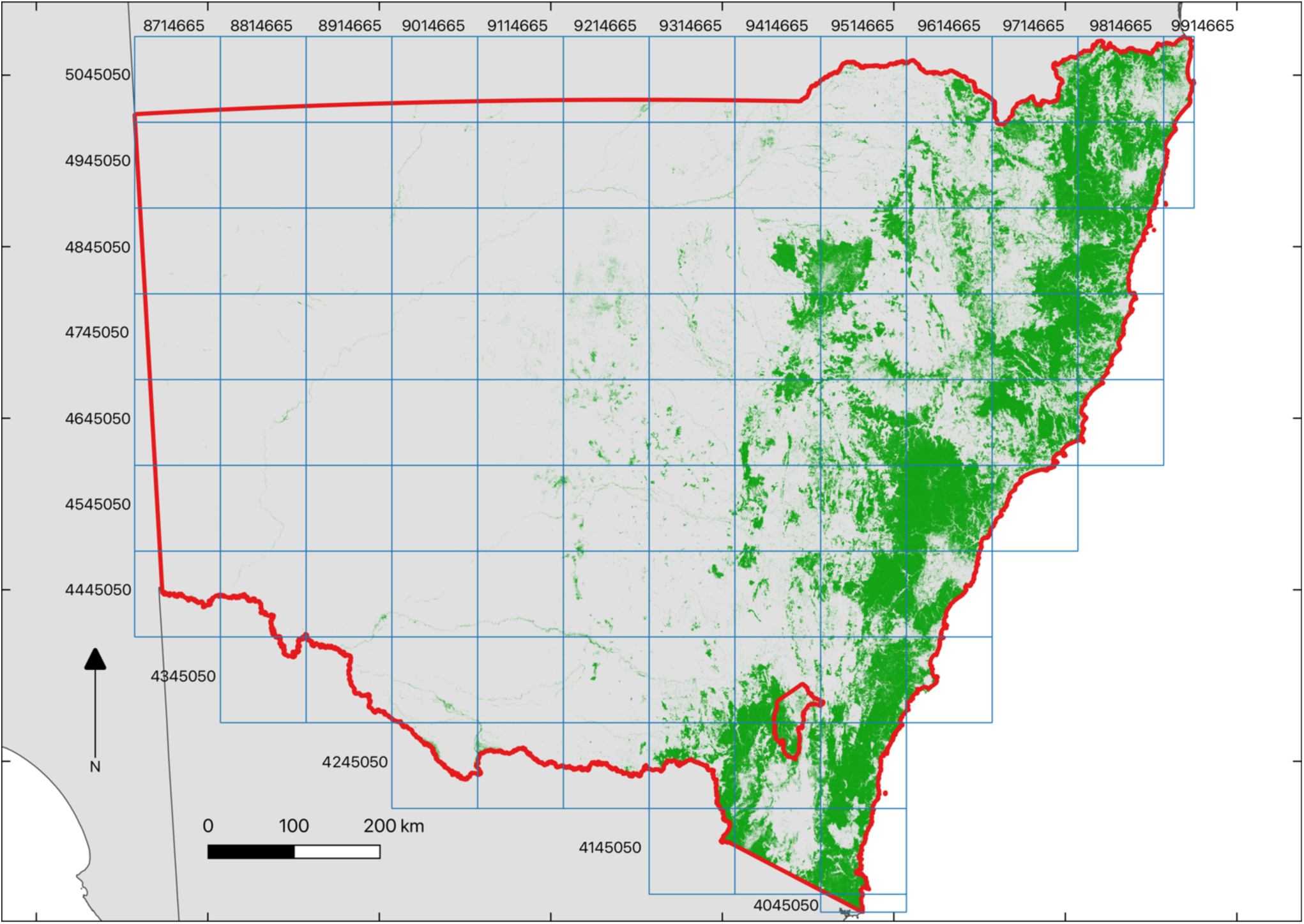
Tiles of the FMC data cube across the states of NSW and the ACT (red borders) in continental Australia (grey area). Tile boundaries are shown in blue with corresponding coordinates shown at the start of columns (x) or rows (y), which represent the top-left coordinates and names of associated tiles. The shaded, green area indicates the forests and woodlands of the FMC dataset.

### F. Variation of FMC in NSW

FMC variation is very different across the state, from one to 53.9% DMC (standard deviation, Fig. S8). It is lowest in parts of the coastal areas of the state, particularly the low to mid slopes of the north coast as well as some upper slopes of that region and the central coast. Higher elevations in the western slopes and plains region have low variation relative to the rest of that region, which also includes very high variation sometimes within relatively close distances (e.g. Coolah Tops). Certain patches of the mountainous regions in the south-east seem exhibitto have low variation, but in neighbouring areas of that region variation is high. The cCoastal hinterland of the very south-east corner has very low variation. The western slopes region, in the centre of the state, contains forests with the greatest variation in FMC.

**Fig. S8.**
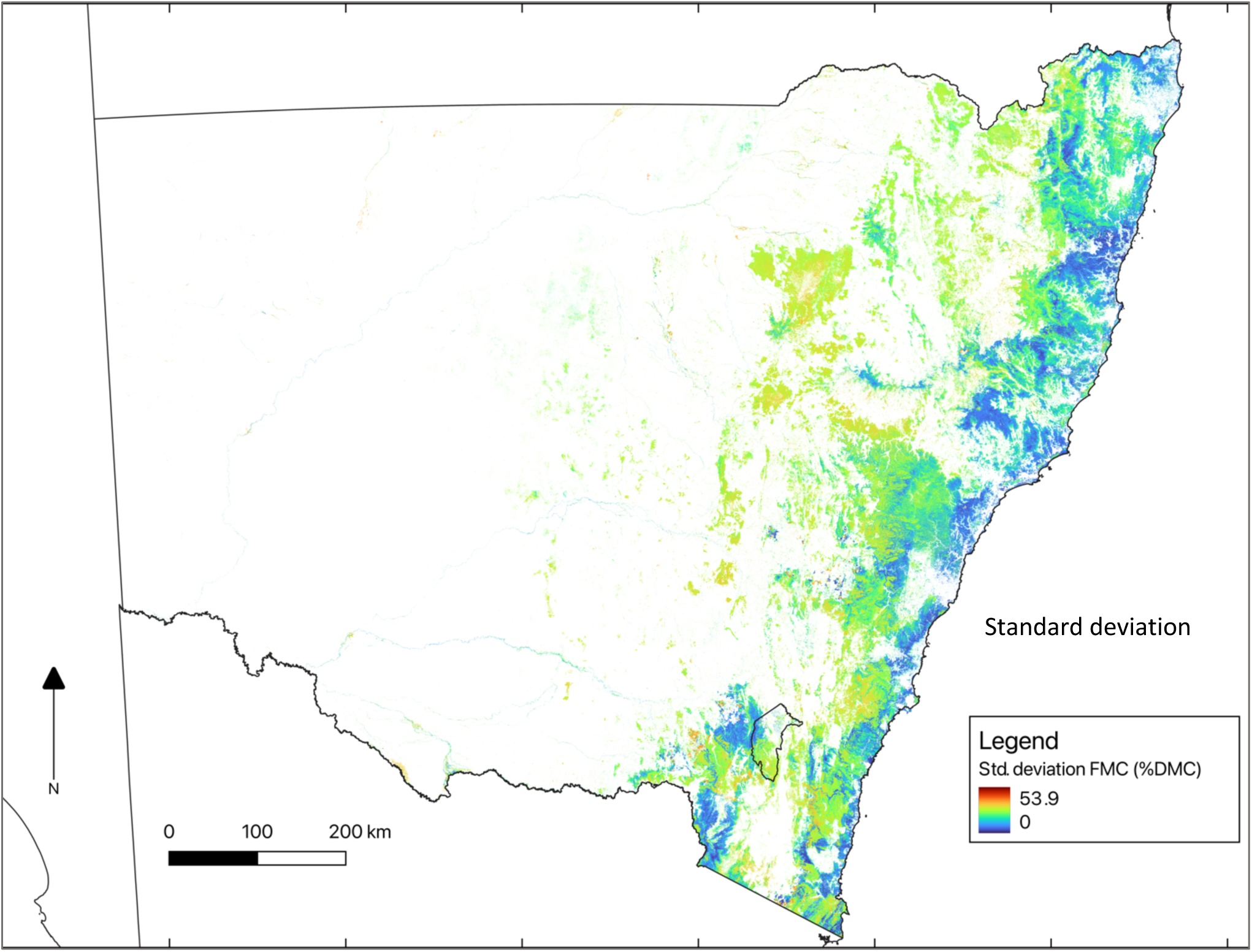
Standard deviation of foliar moisture content (FMC, % dry matter content [DMC]) in forests and woodlands of NSW (2015-2024).

### G. Satellite Bands and Indices Used in RTM Inversion and RF Emulator

**TABLE S1.** Satellite reflectance band used in the RTM inversion in this study (Sentinel-2) and the study of woodland FMC in which the LUT used here was developed (MODIS) [40]. Bold indicates bands used in the RF emulator in this study.

| Band no. | Wavelengths | Used |
| --- | --- | --- |
| **Sentinel-2 (this study)** |  |  |
| 1 | 442 - 453 | False |
| **2** | **491 - 525** | **False** |
| **3** | **559 - 578** | **True** |
| **4** | **664 - 680** | **True** |
| **5** | **703 - 712** | **True** |
| **6** | **740 - 748** | **True** |
| **7** | **782 - 793** | **True** |
| **8** | **832 - 886** | **True** |
| **8a** | **864 - 875** | **True** |
| 9 | 944 - 955 | False |
| 10 | 1373 - 1389 | False |
| **11** | **1613 - 1659** | **True** |
| **12** | **2201 - 2290** | **True** |
| **MODIS (see [40])** |  |  |
| 1 | 620 - 670 | True |
| 2 | 841 - 876 | True |
| 3 | 459 - 479 | True |
| 4 | 545 - 565 | True |
| 5 | 1230 - 1250 | True |
| 6 | 1628 - 1652 | True |
| 7 | 2105 - 2155 | True |

### H. Field Site Details and Species

**TABLE S2.** Field site details for FMC data collection used in remote-sensing model validation. Sites are located in the Central coast, Riverina, slopes and tablelands regions of NSW. Plant community types are described in Roff et al. [50].

| Field site | Latitude, Longitude | Plant community type | Land manager |
| --- | --- | --- | --- |
| **Blue Mountain Ridge** | -33.660198°,<br>150.617703° | *Sydney Hinterland*<br>*Enriched Sandstone*<br>*Bloodwood Forest* | NSW National Parks<br>and Wildlife Service<br>(NPWS) |
| **Bowen Mountain** | -33.5945064°,<br>150.6256786° | *Sydney Hinterland*<br>*Peppermint-Apple*<br>*Forest* | NSW NPWS |
| **Cattai** | -33.566813°,<br>150.472000° | *Sydney Hinterland Grey*<br>*Gum Transition Forest* | NSW NPWS |
| **Colenso Road** | -35.5594678°,<br>144.1869144° | *Black Box - Lignum*<br>*woodland wetland of*<br>*the inner floodplains in*<br>*the semi-arid (warm)*<br>*climate zone* | NSW department<br>lands, reserves and<br>cemeteries |
| **Conargo Road** | -35.358700°,<br>145.502979° | *Black Box - Lignum*<br>*woodland wetland of*<br>*the inner floodplains in*<br>*the semi-arid (warm)*<br>*climate zone* | NSW department<br>lands, reserves and<br>cemeteries |
| **Coreen State Forest** | -35.7151501°,<br>146.3354911° | *Western Grey Box tall*<br>*grassy woodland on*<br>*alluvial loam and clay*<br>*soils in the NSW South*<br>*Western Slopes and*<br>*Riverina Bioregions* | Forestry Corporation of<br>NSW |
| **Doona State Forest** | -31.353108°,<br>150.404037° | *Narrow-leaved Ironbark*<br>*- cypress pine - White*<br>*Box shrubby open forest*<br>*in the Brigalow Belt*<br>*South Bioregion and*<br>*Nandewar Bioregion* | Forestry corporation of<br>NSW |
| **Goobang** | -32.963502°,<br>148.427562° | *White Box grassy woodland in the upper slopes sub-region of the NSW South Western Slopes Bioregion* | NSW NPWS |
| **Goombargana Reserve** | -35.7434852°,<br>146.5603211° | *Yellow Box - White Cypress Pine grassy woodland on deep sandy-loam alluvial soils of the eastern Riverina Bioregion and western NSW South Western Slopes Bioregion* | NSW department lands, reserves and cemeteries |
| **Kentlyn** | -34.079180°,<br>150.846420° | *Sydney Hinterland Grey Gum Transition Forest* | NSW department lands, reserves and cemeteries |
| **Mallan** | -35.1252788°,<br>143.7044434° | *Black Box open woodland wetland with chenopod understorey mainly on the outer floodplains in south-western NSW (mainly Riverina Bioregion and Murray Darling Depression Bioregion)* | NSW department lands, reserves and cemeteries |
| **Moulamein** | -35.107475°,<br>144.032873° | *Black Box - Lignum woodland wetland of the inner floodplains in the semi-arid (warm) climate zone (mainly Riverina Bioregion and Murray Darling Depression Bioregion)* | NSW department lands, reserves and cemeteries |
| **Mount Arthur** | -32.554351°,<br>148.885252° | *White Box grassy*<br>*woodland in the upper*<br>*slopes sub-region of the*<br>*NSW South Western*<br>*Slopes Bioregion* | NSW department<br>lands, reserves and<br>cemeteries |
| **Nangar** | 148.516625°, -<br>33.421945° | *White Box - Black*<br>*Cypress Pine - red gum*<br>*+/- Mugga Ironbark*<br>*shrubby woodland in*<br>*hills of the NSW central*<br>*western slopes* | NSW NPWS |
| **Pilliga** | -31.098686°,<br>149.1392149° | *White Cypress Pine -*<br>*Narrow-leaved Ironbark*<br>*- White Bloodwood -*<br>*red gum shrub grass*<br>*woodland of the Pilliga*<br>*- Coonabarabran*<br>*region, Brigalow Belt*<br>*South Bioregion* | NSW NPWS |
| **River Moulamein** | -35.096342°,<br>144.0295974° | *River Red Gum-sedge*<br>*dominated very tall*<br>*open forest in*<br>*frequently flooded*<br>*forest wetland along*<br>*major rivers and*<br>*floodplains in south-*<br>*western NSW* | NSW department<br>lands, reserves and<br>cemeteries |
| **South West**<br>**Woodlands** | -35.491818°,<br>145.719151° | *Western Grey Box tall*<br>*grassy woodland on*<br>*alluvial loam and clay*<br>*soils in the NSW South*<br>*Western Slopes and*<br>*Riverina Bioregions* | NSW NPWS |
| **The Rock** | -35.262604°,<br>147.074177° | *Dwyers Red Gum - Black Cypress Pine - Currawang shrubby low woodland on rocky hills mainly in the NSW South Western Slopes Bioregion* | NSW NPWS |
| **Trinkey** | -31.40578°,<br>149.994891° | *Narrow-leaved Ironbark - White Cypress Pine - Buloke tall open forest on lower slopes and flats in the Pilliga Scrub and surrounding forests in the central north Brigalow Belt South Bioregion* | NSW NPWS |
| **Walbundrie Culcairn Reserve** | -35.6649884°,<br>146.8967424° | *Western Grey Box tall grassy woodland on alluvial loam and clay soils in the NSW South Western Slopes and Riverina Bioregions* | NSW department lands, reserves and cemeteries |
| **Warumbungles** | -31.293034°,<br>149.039300° | *Inland Scribbly Gum - White Bloodwood - Red Stringybark - Black Cypress Pine shrubby sandstone woodland mainly of the Warrumbungle NP - Pilliga region in the Brigalow Belt South Bioregion* | NSW NPWS |
| **Yanga National Park** | -34.696155°,<br>143.634300° | *Black Box open woodland wetland with chenopod understorey mainly on the outer floodplains in south-western NSW (mainly Riverina Bioregion and Murray Darling Depression Bioregion)* | NSW NPWS |
| **Yanga SCA** | -34.770211°,<br>143.773269° | *Belah/Black Oak - Western Rosewood - Wilga woodland of central NSW including the Cobar Peneplain Bioregion* | NSW NPWS |

**TABLE S3.** Frequency of species collected per sample. Note: validation values represent the average of three samples per site, which explains the high frequency of samples relative to the total number of validation sites.

| **Species collected** | **Sample frequency** |
| --- | --- |
| *Eucalyptus largiflorens* | 30 |
| *Eucalyptus microcarpa* | 24 |
| *Eucalyptus albens* | 18 |
| *Eucalyptus crebra* | 8 |
| *Eucalyptus punctata* | 8 |
| *Eucalyptus fibrosa* | 7 |
| *Eucalyptus rossii* | 6 |
| *Eucalyptus camaldulensis* | 6 |
| *Eucalyptus dwyeri*, *E. blakelyi* | 6 |
| *Eucalyptus punctata* | 5 |
| *Eucalyptus piperita/Eucalyptus consideniana* | 4 |
| *Corymbia gummifera* | 4 |
| *Eucalyptus macrorhyncha* | 4 |
| *Angophora costata* | 3 |
| *Corymbia sp.* | 3 |
| *Eucalyptus eugenioides* | 2 |
| *Eucalyptus sideroxylon* | 2 |
| *Eucalyptus globoidea* | 2 |
| *Eucalyptus eugenioides* | 1 |

**TABLE S4.** Species of trees at each field site sampled for FMC estimates to validate the satellite-based model.

| Site ID | Site name | Species sampled |
| --- | --- | --- |
| 1 | Blue Mountain Ridge | *Eucalyptus globoidea*, *Corymbia gummifera*, *Angophora costata* |
| 2 | Bowen Mountain | *Eucalyptus piperita/Eucalyptus consideniana*, *C. gummifera* |
| 3 | Cattai | *Eucalyptus punctata*, *A. costata* |
| 4 | Colenso Road | *Eucalyptus largiflorens* |
| 5 | Conargo Road | *Eucalyptus microcarpa* |
| 6 | Coreen State Forest | *E. microcarpa* |
| 7 | Doona State Forest | *Eucalyptus crebra* |
| 8 | Goobang | *Eucalyptus albens* |
| 9 | Goombargana Reserve | *E. albens* |
| 10 | Kentlyn | *Eucalyptus eugenioides*, *E. punctata*, *Corymbia gummifera* |
| 11 | Mallan | *E. largiflorens* |
| 12 | Moulamein | *E. largiflorens* |
| 13 | Mount Arthur | *E. albens* |
| 14 | Nangar | *Eucalyptus sideroxylon*, *Eucalyptus macrorhyncha* |
| 15 | Pilliga | *Eucalyptus fibrosa*, *Eucalyptus crebra* |
| 16 | River Moulamein | *Eucalyptus camaldulensis* |
| 17 | South West Woodlands | *E. microcarpa* |
| 18 | The Rock | *Eucalyptus dweryi*, *E. blakelyi* |
| **19** | Trinkey | *E. fibrosa* |
| **20** | Walbundrie Culcairn Reserve | *E. microcarpa* |
| **21** | Warumbungles | *Eucalytpus rossii* |
| **22** | Yanga National Park | *E. largiflorens* |
| **23** | Yanga SCA | *E. largiflorens* |

